# Basal forebrain and brainstem cholinergic neurons differentially impact amygdala circuits and learning-related behavior

**DOI:** 10.1101/368134

**Authors:** Teemu Aitta-aho, Y Audrey Hay, Benjamin U. Phillips, Lisa M. Saksida, Tim J. Bussey, Ole Paulsen, John Apergis-Schoute

## Abstract

The central cholinergic system and the amygdala are important for motivation and mnemonic processes. Different cholinergic populations innervate the amygdala but it is unclear how these projections impact amygdala processes. Using optogenetic circuit-mapping strategies in ChAT-cre mice we demonstrate that amygdala-projecting basal forebrain and brainstem ChAT-containing neurons can differentially affect amygdala circuits and behavior. Photo-activating ChAT terminals *in vitro* revealed the underlying synaptic impact of brainstem inputs to the central lateral division to be excitatory, mediated via the synergistic glutamatergic activation of AMPA and NMDA receptors. In contrast, stimulating basal forebrain inputs to the basal nucleus resulted in endogenous ACh release resulting in biphasic inhibition-excitation responses onto principal neurons. Such response profiles are physiological hallmarks of neural oscillations and could thus form the basis of acetylcholine-mediated rhythmicity in amygdala networks. Consistent with this, *in vivo* NBm activation strengthened amygdala basal nucleus theta and gamma frequency rhythmicity, both of which continued for seconds after stimulation and were dependent on local muscarinic or nicotinic receptor activation, respectively. Activation of brainstem ChAT-containing neurons however resulted in a transient increase in central lateral amygdala activity that was independent of cholinergic receptors. In addition, driving these respective inputs in behaving animals induced opposing appetitive and defensive learning-related behavioral changes. Since learning and memory is supported by both cellular and network-level processes in central cholinergic and amygdala networks, these results provide a route by which distinct cholinergic inputs can convey salient information to the amygdala and promote associative biophysical changes that underlie emotional memories.

## Introduction

Effective learning requires attending selectively to information while ignoring distractions [1]. The capacity of a stimulus to capture attention can be attributed to the stimulus’ saliency or features that make it stand out among other stimuli [2]. Both central cholinergic systems [4,5]and the amygdala [5,6] are important for saliency detection and attention – processes that are critical for effective learning and memory [7–9] that when disrupted can result in diseases that severely affect memory. Alzheimer’s disease patients, for example, show severe impairments in emotional memory [10–12] – a phenotype that may result from a significant reduction in amygdala and central cholinergic cell populations [11,13,14]. Such relations between circuit-level degeneration and disturbances in cognition underscore the clinical significance of acetylcholine (ACh) and amygdala network interactions.

Mechanistically, central ACh can strengthen attention by enhancing the gain of sensory signaling [15–18]. Moreover, by synchronizing activity between neural networks[19–21], ACh can support attentional [22,23] and mnemonic [7] processes and promote synaptic plasticity [24,25]. Cholinergic populations in the basal forebrain (NBm) and the pedunculopontine tegmental nucleus (PPT) of the brainstem show overlapping functions in regards to attention and motivation [26,27]. Moreover, individual NBm neurons have been shown to respond to natural and learned emotionally-salient stimuli [28–30] and can do so on the order of tens of milliseconds [31]. These results suggest that NBm neurons are positioned to quickly communicate value-based information to target regions. Brainstem neurons similarly respond to salient cues and transiently discharge in response to reward delivery [32].

Both cholinergic NBm [33] and PPT populations [34,35] innervate the amygdala, a collection of limbic nuclei that cooperate in forming associations between neutral environmental stimuli and those with intrinsic value [36]. Similar to cholinergic neurons, ones in the basolateral nucleus (BL) also encode rewarding and aversive stimuli [37,38] and can update their activity when stimulus valence is modified [38,39]. Despite these anatomical and functional relationships the impact of distinct cholinergic populations on amygdala processes has not been thoroughly compared and contrasted. Moreover, it is unclear whether ACh inputs to the amygdala can support rhythmic neural activity as they do in other memory-related brain regions. Increased synchrony in amygdala circuits has been linked to innate [40,41] and learned emotional responses [42,43] but the mechanisms mediating this rhythmicity are currently unresolved.

To shed light on the cholinergic influences on amygdala function this study has implemented optogenetic circuit mapping methods in transgenic ChAT-cre mice to determine the functional relationship between cholinergic inputs to the amygdala and amygdala-dependent learning and memory. We show that cholinergic NBm populations that target the BL can synchronize ongoing BL Aitta-aho et al. Cholinergic control of amygdala function activity through cholinergic transmission while brainstem ACh populations can transiently excite central lateral (CeL) amygdala circuits via glutamatergic neurotransmission. Behaviorally, photo-activating BL- and CeL-projecting ChAT pathways induced opposing appetitive and defensive learning-related behavioral responses. Collectively, these results highlight the unique role of distinct central cholinergic systems on amygdala circuits and function.

## Results

### Non-overlapping innervation of amygdala nuclei by distinct cholinergic cell populations

Cholinergic fibers emanating from the NBm [33] and PPT [34,35] project to the amygdala but their functional impact has not been thoroughly delineated. To investigate these projections we used a cre-recombinase approach in transgenic ChAT-cre mice to express the light-activated sodium channel channelrhodopsin-GFP (ChR2) in NBm and PPT cholinergic populations and their axons (Fig 1). Expression analysis revealed that GFP-labeled NBm and PPT neurons were mostly immuno-positive for the ACh-related enzyme ChAT (NBm: n = 3 mice, 96.4 ± 1.3%, 172 cells; PPT: n = 4 mice, 89.2 ± 4.9% 117 cells) (Fig S1). Axons from NBm-ChAT neurons densely innervated the BL and central medial (CeM) divisions of the amygdala leaving the lateral (LA) and CeL nuclei remarkably devoid of NBm-ACh inputs (Fig 1B,C). In contrast, PPT-ChAT neurons almost exclusively innervated the CeL nucleus (Fig 1E,F). These anatomical results reveal a striking non-overlapping and complementary innervation of anatomically and functionally distinct amygdala nuclei by different ACh populations.

**Figure 1:**
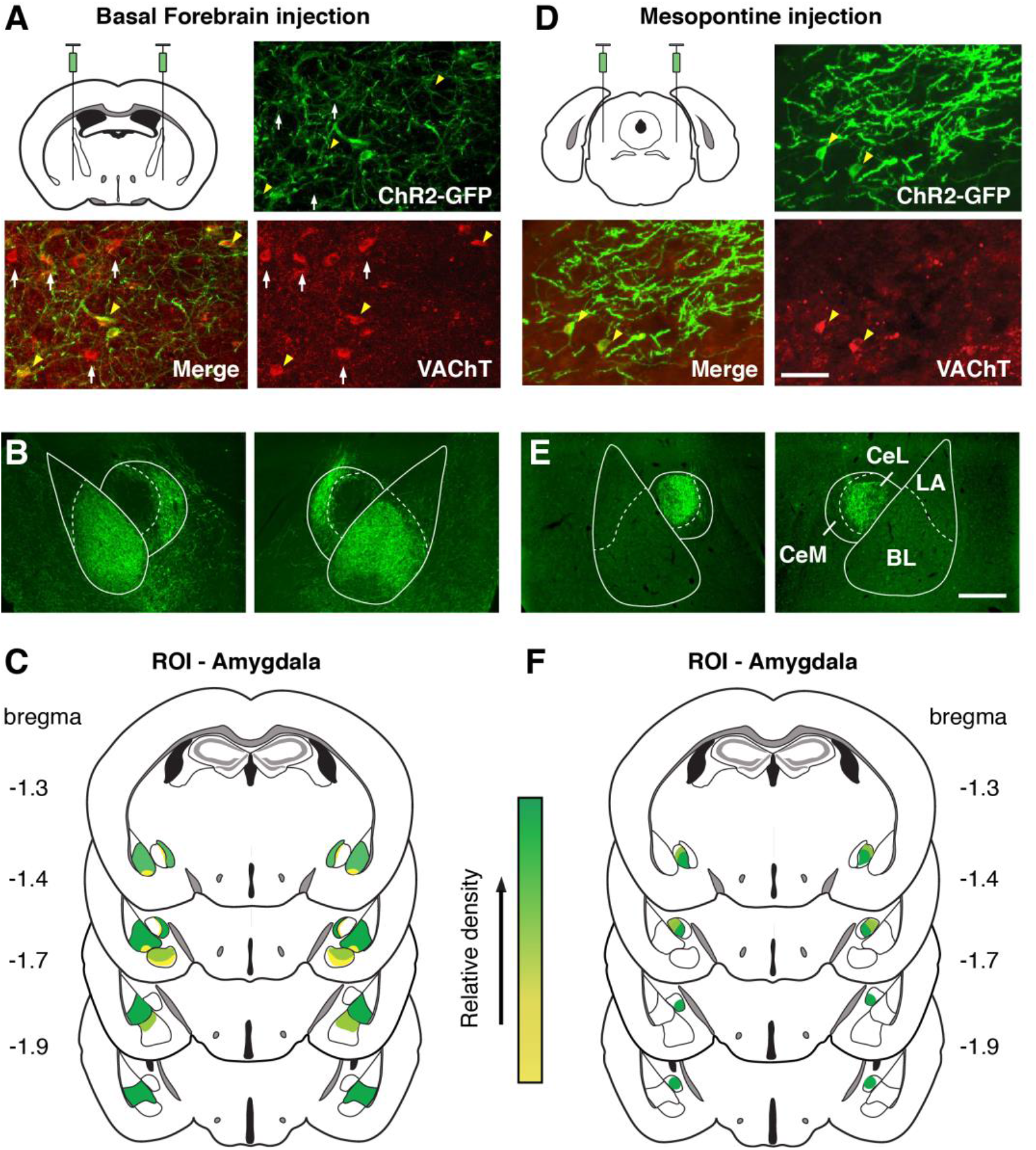
Basal forebrain and Brainstem cholinergic neurons differentially innervate the amygdala. Viral infection of NBm (A) and PPT (D) ChAT-containing neurons in transgenic ChAT-cre mice led to ChR2-GFP expression in local VAChT-expressing neurons (A,D). GFP-labeled fibers of NBm-ChAT neurons mostly innervated the basal and central medial division of the amygdala (B,C) while PPT-ChAT neurons sent projections to the central lateral amygdala nucleus (E,F). C,F. Group summary of BF (C) and PPT (F) distribution in the amygdala plotted as relative density by color. Abbreviations: LA, lateral amygdala; BL, basolateral amygdala; CeL, central lateral amygdala; CeM, central medial amygdala. ROI, region of interest. Scale bars: A, D, 100 μm; B, E, 1 mm.

### *In vivo* photo-activation of NBm-ChAT neurons synchronizes BL activity for seconds after stimulation while PPT-ChAT neurons transiently activate CeL networks

Next, we characterized the impact of NBm- and PPT-ChAT stimulation on BL and CeL network activity. Local field potentials (LFP) and multi-unit activity (MUA) were recorded from the NBm and BL or PPT and CeL in urethane anesthetized ChAT-cre mice expressing ChR2 in ChAT neurons. As expected, 20Hz photo-activation of NBm-ChAT circuits increased the power intensity of the local LFP profoundly in the slow gamma range (20–30Hz) (Fig 2A,B, blue trace) as well as more broadly in theta (4–16Hz) and extended gamma (20–60Hz) that decayed following stimulation (Fig 2A,B; red trace). Simultaneous BL-LFP recordings showed a marked increase of theta and slow gamma (Fig 2C,D; blue trace) that, in contrast to NBm responses, remarkably persisted and increased in power after stimulation had ended (Fig 2C,D; red trace). Similar NBm-ChAT-evoked BL responses were recorded in awake mice (n = 4) albeit of a smaller magnitude (Fig S2).

**Figure 2:**
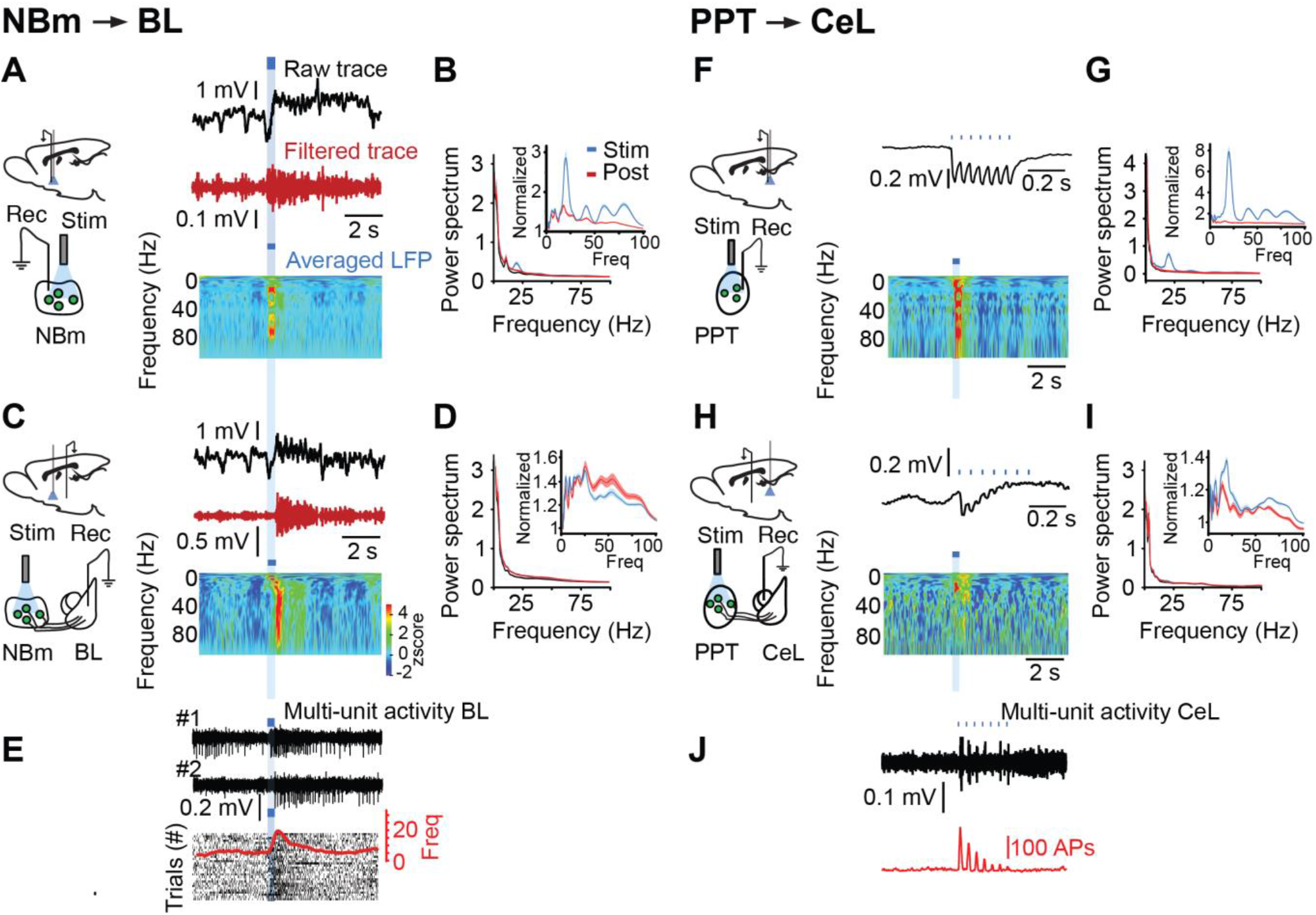
In vivo PPT/NBm photo-activation respectively synchronizes local CeL and BL activity transiently and for seconds after stimulation. NBm-ACh stimulation (n = 10) results in an immediate (A,B) and delayed (C,D) change in NBm and BL local field potentials (LFP), respectively. A,C. Top – right, raw unfiltered LFP; Middle, high-pass filtered LFP; Bottom, color-coded wavelet analysis plotting the frequency power over time (A, NBm; C, BL). Power spectrum of NBm (B) and BL (D) LFP during (blue trace) and immediately after (red trace) stimulation. Normalized NBm (B-inset) and BL (D-inset) power spectrum analysis where the power during (blue trace) and after (red trace) stimulation was divided by the baseline power spectrum. E. (Top) Two separate examples of multi-unit BL activity in response to NBm photo-stimulation. (Bottom) Rasterplot of multi-unit activity recorded during multiple sweeps and their average response (red trace). B-J. Similar analyses for PPT photo-induced responses in the PPT and CeL (n = 6). Photo-driven responses and power spectrum of PPT (F,G) and CeL (H,I) activity. J. PPT photo-evoked multi-unit activity from CeL recordings. Power Spectrum units, mV^2^/Hz (B,D; G,I).

In response to PPT-ChAT activation, PPT (Fig 2F) and CeL (Fig 2H) circuits showed time-locked responses to the photo-activation frequency (20Hz) that was greater during the stimulation than after (Fig 2D, 2I) revealing, in comparison to NBm-ChAT projections to the BL, a more temporally restricted influence of PPT-ChAT projections on CeL networks. Both NBm- and PPT-ChAT evoked LFPs were likely due to local changes in amygdala activity since stimulation increased BL (Fig 2E) and CeL (Fig 2J) MUA. Together, these results indicate that 20 Hz photo-activation of NBm-ChAT circuits *in vivo* can effectively synchronize ongoing BL network activity for seconds after stimulation while PPT activation transiently influences CeL neurons.

### NBm-ChAT neurons evoke biphasic cholinergic responses in BL networks while PPT-ChAT inputs excite CeL neurons through AMPA and NMDA receptor activation

We next investigated in acute brain slices the cellular mechanisms underlying NBm- (Fig 3A-H) and PPT-ChAT (Fig 3I-N) mediated responses in BL and CeL neurons. Consistent with previous work [44,45], current-clamp recordings from BL principal cells showed that minimal light stimulation (0.5 ms) of NBm-ChAT inputs to the BL can effectively suppress amygdala activity (n = 29 out 41 cells, Fig 3D). These responses persisted with glutamate (CNQX, AP-V 20 μM; AP-V, 100 μM)) and GABA_A,B_ (gabazine 10 μM; CGP-52432 10 μM) receptor antagonists present (Fig S3A) and were of similar latencies to control conditions (n = 11 cells, 43.3 ± 2.5 and 40.7 ± 3.1 ms, Wilcoxon Signed Rank Test, p = 0.65). These inhibitory responses reversed at -90 mV (n=6) (Fig S2A), were insensitive to the SK channel blocker apamin (% of baseline, 96.1 ± 3.2%; n = 5; Wilcoxon Signed Rank Test, p = 0.38; Fig S2B) but were abolished by application of the muscarinic M1 receptor (M1R)-antagonist pirenzepine (PRZ) (n = 8 cells: 1 μM; Amplitude: ctl, 9.4 ± 1.6, PRZ, 1.4 ± 0.3 pA; Wilcoxon Signed Rank Test; p =0.001; Fig 3E; S3C) indicating that they were due to changes in K^+^ conductances.

**Figure 3:**
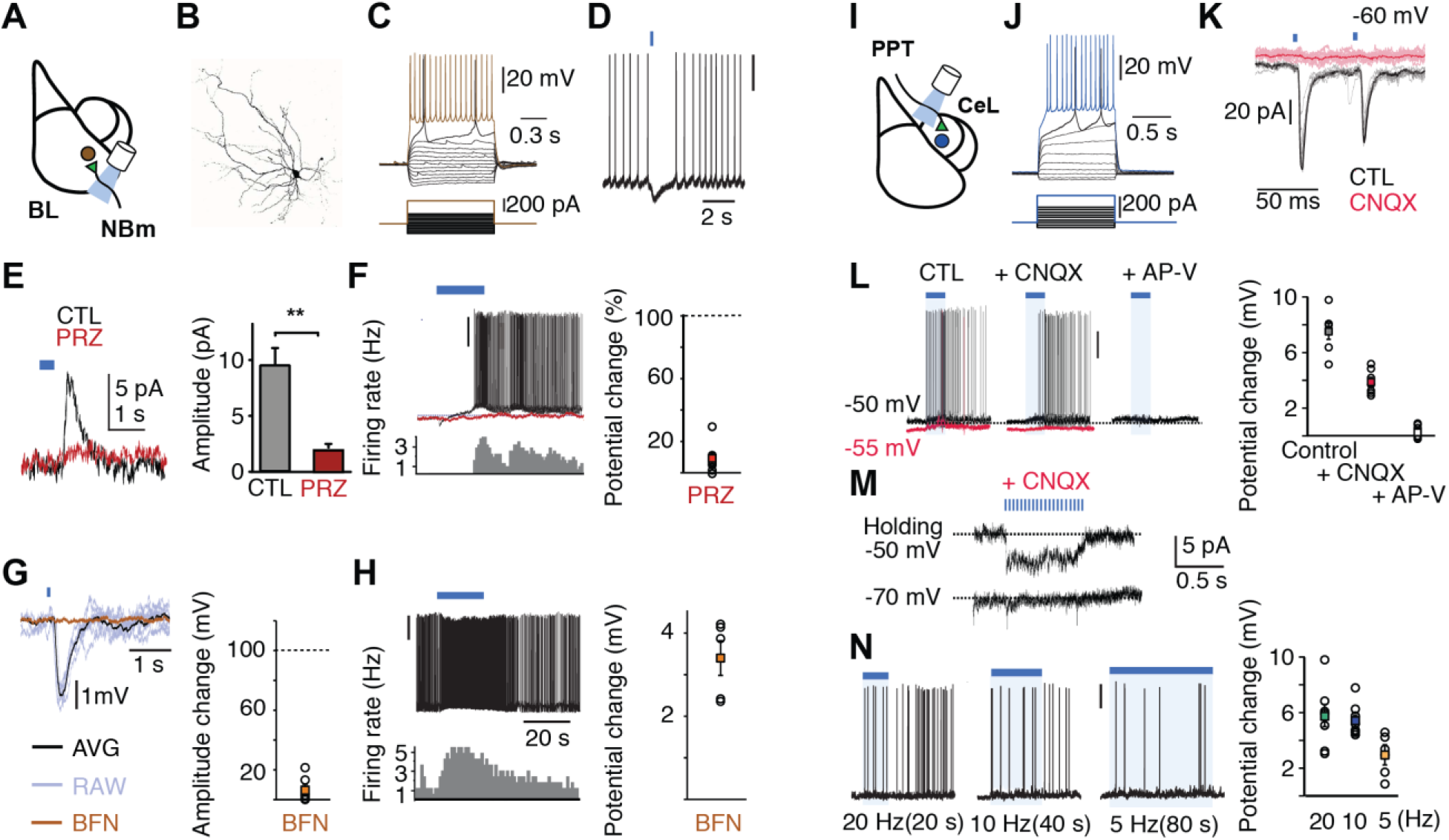
Electrophysiological impact of NBm and PPT cholinergic populations onto BL (A) and CeL (I) activity *in vitro*. A-H. Activation of NBm to BL inputs results in a muscarinic M1 receptor-dependent fast inhibition followed by slow excitation. Morphology (B) Current-voltage relationship (C) and of a BL principal neuron. All recordings showed similar electrophysiological signatures. Optogenetic terminal activation with 1 ms flash of blue light suppressed principal neuron action potentials (D) by increasing a PRZ-sensitive outward K+ conductance (E). In the presence of synaptic blockers, 20 Hz NBm-BL photo-stimulation (20 Hz for 20 sec) led to a PRZ-sensitive slow membrane depolarization in BL principal neurons (F). Occlusion of inwardly-rectifying K^+^ channels with the GABA_b_ agonist baclofen (20 μM) with glutamate and GABA receptors antagonists present blocked the NBm-BL evoked fast IPSC (2.1 ± 4.0% of control response, n=6 cells, Wilcoxon Signed Rank Test, p = 0.04) (G). With baclofen present, 20 Hz NBm-BL photo-stimulation increased BL neuronal firing by depolarizing (3.4 ± 0.4 mV) BL principal neurons (H). I-N, PPT axons directly excite CeL neurons by releasing glutamate that acts postsynaptically on AMPA and NMDA receptors. J, example current-voltage relationship of recorded CeL neuron. K, brief blue light pulses evoked post-synaptic currents that were sensitive to the AMPA receptor antagonist CNQX (20 μM). Stimulation-induced increase in CeL firing frequency (L-left) is composed of an AMPA-mediated (L-middle) and NMDA-mediated (L-right) depolarization, the NMDA response being absent at a more negative V_m_ (M). N, stimulation at 20, 10, and 5 Hz was sufficient to excite CeL neurons.

The BL contains a relatively high degree of acetylcholinesterase (AChE), the enzyme that catalyzes the breakdown of ACh, suggesting that BL cholinergic transmission is tightly regulated both spatially and temporally. It is thus conceivable that the NBm-ChAT mediated BL inhibition may result from the local release of ACh. To test this hypothesis we evoked inhibitory responses in the presence of synaptic antagonists while bath-applying the AChE inhibitor physostigmine (Phys) (10 μM) (Fig S3C). Blocking ACh breakdown led to an increase in the amplitude and duration of NBm-ChAT evoked inhibition (n = 6 cells: Amplitude: ctl, 9.6 ± 1.6; physo, 25.9 ± 6.1 pA. Duration: ctl, 0.08 ± 0.02; physo, 1.23 ± 0.34 s) (Fig 3B3,4). This inhibition were completely blocked by the M1R antagonist PRZ (1 μM) (Fig 3B3,4) (Amplitude: ctl, 9.6 ± 1.6; PRZ, 1.9 ± 0.5; washout, 6.2 ± 1.2 pA. Duration: ctl, 0.08 ± 0.02; PRZ, 0.03 ± 0.01; washout 0.07 ± 0.03 s) (Amplitude: One-way ANOVA; F(3,18) = 9.4 p = 0.0005, Bonferroni post-hoc, Control vs. physo p = 0.0008, Control vs. PRZ p = 0.01, physo vs. PRZ/washout p = 0.004 PRZ vs. washout p = 0.01. Decay Time: One-way ANOVA; F(3,18) = 2.28 p = 0.0001, Bonferroni post-hoc, Control vs. physo p = 0.0006, Control vs. PRZ p = 0.01, physo vs. PRZ/washout p = 0.0002 PRZ vs. washout p = 0.01). These results are in line with previous studies [46,47] and overall demonstrate that ACh is reliably released and acts relatively quickly to robustly inhibit BL principal neurons.

High frequency (20Hz) stimulation of NBm-ChAT to BL inputs often evoked a large inhibitory response followed by a sustained depolarization (Fig S2D, 3F – black trace), that was sensitive to PRZ (7.3 ± 1.4% of control, n = 8 cells, WilcoxonSignedRank-Test, p = 0.008; Fig 3F – red trace). To test whether the M1R-mediated depolarization was independent of the preceding inhibition we used an occlusion strategy, which consisted of maximally activating M1R-coupled K^+^ channels with baclofen (BFN), a GABA_b_ receptor agonist. Photo-stimulation with BFN present eliminated the inhibitory response but left the excitatory response intact (Fig 3H), indicating that the NBm-ChAT mediated excitation is independent of the preceding inhibitory signal.

In contrast to NBm-BL pathways, single CeL light pulses onto PPT-ChAT (Fig 3I) fibers elicited short-latency inward currents in voltage-clamped CeL neurons (n = 19 cells: Amplitude: 24.3 ± 4.5 pA, Latency: 4.2 ± 0.2 ms). Photo-evoked currents were abolished by AMPA receptor (AMPA-R) antagonism (CNQX, 20 μM) (Fig 3K), suggesting that PPT-ChAT inputs to the CeL were glutamatergic. Such a mechanism is not extraordinary since cholinergic populations can communicate via glutamatergic transmission [48,49]. Photo-activating at 20Hz increased CeL activity that outlasted the stimulation duration (Fig 3L-left). While the initial depolarization was dependent on AMPA-R activation (Fig 3L-middle), the sustained increase in excitability was sensitive to the NMDA receptor (NMDA-R) antagonist AP-V (100 μM) (Fig 3L-right) (n = 7 cells: ctl, 7.4 ± 0.5; CNQX, 3.9 ± 0.3; AP-V, 0.5 ± 0.2 mV; One-way ANOVA F(1.57, 9.43) = 128.9, p = 0.0001; Bonferroni post-hoc, Control vs. cnqx p = 0.0009, Control vs. AP-V p = 0.0001, CNQX vs. AP-V p = 0.0007). NMDA responses were mostly absent at hyperpolarized membrane potentials (V_m_) (Fig 3M), suggesting for cooperative interactions between AMPA-R and NMDA-R at depolarized V_m_.

Since PPT neurons fire at various frequencies *in vivo* [50] we next tested different PPT-ChAT photo-stimulation frequencies on CeL neurons (Fig 3N). Photo-stimulation at 10 and 20Hz excite CeL neurons similarly while 5 Hz did so less (n = 8 cells: 20 Hz, 5.7 ± 0.7 mV, n = 8; 10 Hz, 5.3 ± 0.4 mV, n = 8; 5 Hz, 2.9 ± 0.7 mV, n = 5; Two-way ANOVA F(2, 19) = 6.93, p = 0.005; Bonferroni post-hoc, 20Hz vs. 10Hz p = 0.82, 20Hz vs. 5Hz p = 0.01, 10Hz vs. 5Hz p = 0.02). These responses were insensitive to muscarinic (atropine (ATR), 2 μM, 98.1 ± 6.34% of control, n = 5 cells; WilcoxonSignedRank-Test, p = 0.57) or nicotinic (mecamylamine (MEC), 10 μM, 104.3 ± 3.71% of control, n = 8 cells; WilcoxonSignedRank-Test, p = 0.36) receptor antagonists (Fig S3E-G). Moreover, bath-applied ACh (100 μM) did not impact the V_m_ of CeL neurons (baseline, -75.1 ± 2.4; ACh, -73.6 ± 2.5 mV, n = 8 cells, WilcoxonSignedRank-Test, p = 0.11) (Fig S3H,I). These findings provide evidence that PPT-ChAT neurons excite CeL neurons through ACh-independent glutamate release *in vitro*.

### NBm-ChAT neurons increase BL network rhythmicity through acetylcholine receptor activation *in vivo* while PPT-ChAT evoked CeL activity is independent of cholinergic transmission

The above results demonstrate that NBm- and PPT-ChAT have strikingly different mechanisms of action on BL and CeL networks. It remains unclear however whether direct activation of muscarinic receptor is the main mode of activation of BL network *in vivo*. We therefore tested the effects of ACh receptor antagonists on NBm-ChAT evoked BL activity *in vivo* (Fig S5). Stimulations/recordings were made before and after i.p. injections of 4 mg/kg Scopolamine (SCP) and 2 mg/kg MEC. Injections did not significantly affect local NBm 20 Hz (n = 6 mice: Ctl, 736 ± 16; MEC + SCP, 783 ± 18 μV^2^/Hz; Wilcoxon signed-rank test, p = 0.22) (Fig S5B) but significantly reduced NBm-ChAT evoked rhythmicity in the BL (theta; n = 6 mice: Ctl, 1.29 ± 0.03; MEC+SCP, 1.03 ± 0.01 of baseline: slow gamma; Ctl, 1.74 ± 0.04; MEC+SCP, 1.12 ± 0.02 of baseline: fast gamma; Ctl, 1.51 ± 0.04; MEC+SCP, 1.09 ± 0.02 of baseline; WilcoxonSignedRank-Test, p = 0.03 for all comparisons) (Fig S5D).

We next repeated these experiments but instead with BL infusions of ACh-receptor antagonists (Fig 4). Local BL infusions of ATR (0.7 μL) disrupted NBm-ChAT evoked BL-theta specifically following the stimulation (n = 6 mice: Ctl, 1.46 ± 0.10; ATR, 1.12 ± 0.03 of baseline; WilcoxonSignedRank-Test, p = 0.03) (Fig 4E,F) while having no effect on the 20 Hz responses in the NBm (n = 5 mice; Ctl, 715.6 ± 236.6; ATR, 737.6 ± 270.6 μV^2^/Hz); WilcoxonSignedRank-Test, p = 0.76) (Fig 4B). In contrast, BL infusions of MEC (0.02 ng) did not impact theta (n = 6 mice: stim; Ctl, 1.45 ± 0.10, MEC, 1.28 ± 0.10 of baseline, WilcoxonSignedRank-Test p = 0.22: post-stim; ctl, 1.48 ± 0.16 of baseline; MEC, 1.31 ± 0.12, WilcoxonSignedRank-Test p = 0.56) (Fig 4G-,H-left) but significantly affected evoked slow and fast gamma (n = 6 mice: Slow Gamma; stim; Ctl, 1.36 ± 0.11, MEC 1.14 ± 0.05 of baseline, WilcoxonSignedRank-Test p = 0.03: Fast Gamma; stim; Ctl, 1.25 ± 0.10, MEC 1.11 ± 0.05 of baseline, WilcoxonSignedRank-Test p = 0.03) (Fig 4G,H-middle, right) while not affecting NBM-20 Hz responses (n = 6 mice; Ctl, 1026.37 ± 340.52; MEC, 902.56 ± 310.82 μV^2^/Hz); WilcoxonSignedRank-Test, p = 0.06) (Fig 4B). These results indicate that ACh released from NBm terminals can coordinate ongoing BL theta and gamma activity through independent muscarinic and nicotinic receptor activation.

**Figure 4:**
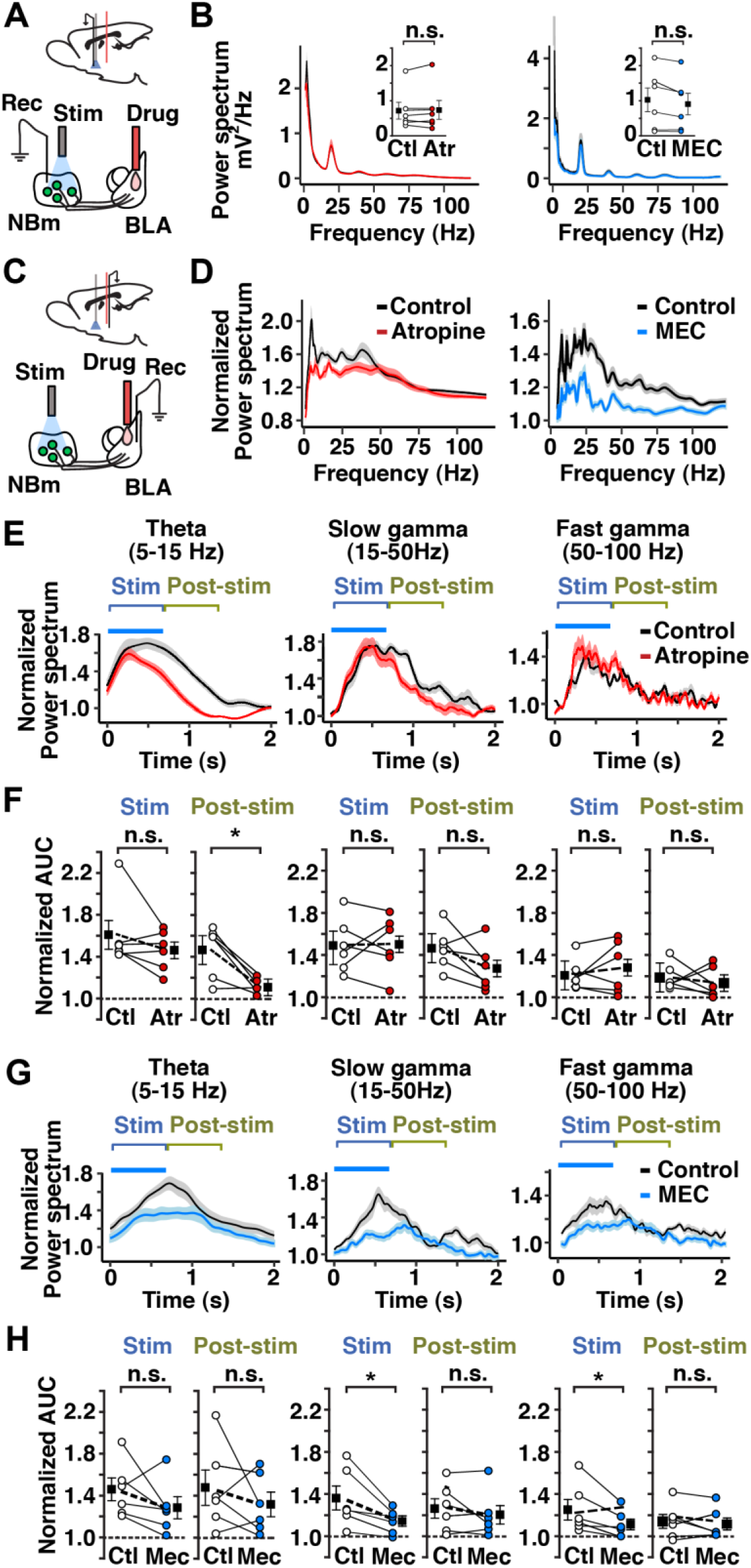
Sustained BL theta rhythmicity in response to NBm-ChAT photo-stimuation is dependent on muscarinic receptor activation while nicotinic receptors contribute to stimulation-induced gamma. A. BL infusions on the muscarinic and nicotinic receptor antagonists, have no significant impact on the magnitude of NBm field potentials (B). BL infusions of ATR (C) does not significantly alter theta, slow and fast gamma power during stimulation but selectively reduces the magnitude of the ongoing NBm-ChAT evoked BL theta activity (D-left; E,F – left). In contrast, BL infusions of MEC significantly reduces NBm-ChAT evoked slow and fast gamma during stimulation(G, H-middle, right). Power Spectrum units(B), μV^2^/Hz.

In contrast to the NBm-BL pathway, CeL responses evoked by PPT-ChAT photo-stimulation were unaffected by ACh receptor antagonism (n = 7 mice: 20 Hz; Ctl, 1.30 ± 0.04; MEC+SCP, 1.26 ± 0.06 of baseline; WilcoxonSignedRank-Test, p = 0.57) (Fig S6C-F). These results are consistent with the *in vitro* results demonstrating that PPT-evoked changes in CeL activity resulted were due to AMPA and NMDA receptor activation (Fig 3I-N). Worthwhile to note is that despite the lack of cholinergic influence of CeL circuits, i.p. injections of MEC + SCP significantly reduced 20 Hz power locally in the PPT (n=7 mice: Ctl, 880 ± 103; MEC+SCP, 1785 ± 101 μV^2^/Hz; WilcoxonSignedRank-Test, p = 0.04) (Fig S6B,right) suggesting for activation of a subpopulation of locally acting PPT-ACh neurons or efferent-specific modes of transmission for PPT-ChAT circuits similar to a subpopulation of dopamine neurons where glutamate and dopamine from the same axon are released from distinct terminal sites [52].

### Distinct ChAT-containing populations reinforce amygdala-dependent approach and avoidance behavior

Prior studies have shown that CeL and accumbal-projecting BL neurons respond preferentially [52] to aversive and appetitive stimuli, respectively, which is consistent with circuit-mapping studies showing that photo-stimulating BL inputs to the CeL or nucleus accumbens result in defensive [53] and approach [53,54] behavior. Moreover, different amygdala populations respond to appetitive and aversive stimuli [37,38] but it is unclear which inputs to the amygdala can communicate such information. We therefore next tested whether NBm-BL and PPT-CeL ChAT projections can influence appetitive and/or aversive amygdala pathways. For this we utilized a conditioned place-preference paradigm where mice were free to explore the entire chamber and photo-activated PPT-CeL or NBm-BL ChAT axons only when mice were within the boundaries of a distinct region (Fig 5A,D). Baseline testing revealed that neither control nor ChR2 mice preferred either region (Habituation: GFP (n = 9 mice), 60.2 ± 6.6%; ChR2, (n = 11 mice) 55.6 ± 5.8%) (Fig 5E,F). Region-specific stimulation however resulted in PPT-CeL ChR2 mice spending less time than GFP-only control mice in the stimulation-paired region (Conditioning: GFP, 55.1 ± 7.5; ChR2, 34.3 ± 6.1%) (Fig 5D,E). Remarkably, after a single conditioning session, ChR2 mice continued to avoid the stimulation-paired region during a “no-stimulation” test phase (Testing; GFP 56.4 ± 8.5, ChR2 33.9 ± 5.8%) (Two-Way ANOVA Group F(2,36) = 4.1 p = 0.02; Bonferonni post-hoc; Hab, Cond, Test - GFP vs ChR2, p = 0.72, p = 0.02, p = 0.01, respectively: Two-Way ANOVA Time F(2,36) = 5.1 p = 0.01; Bonferonni post-hoc; ChR2, Hab vs Cond p = 0.01; Hab vs Test p = 0.01) (Fig 5D,E). Compared to baseline, during both conditioning and testing phases ChR2 mice spent less time in the stimulation-paired region (Normalized data: Cond/Hab GFP = 1.02 ± 0.13, ChR2 = 0.63 ± 0.12, Unpaired T-Test p = 0.03; Test/Hab GFP = 1.05 ± 0.19, ChR2 = 0.61 ± 0.12, Unpaired T-Test p = 0.043) (Fig 5F). Interestingly, during conditioning and testing phases ChR2 mice were slower when in the stimulation-paired area (Two-way ANOVA Group F(1,18) = 7.0 p = 0.01; Bonferonni post-hoc Hab, Cond, Test – GFP vs ChR2, p = 0.58, p = 0.008, p = 0.02: Time F(2,36) = 25.9) (Normalized data in Photostim area: Cond/Hab; GFP 0.93 ± 0.13, ChR2 0.43 ± 0.06, Unpaired T-Test p = 0.01; Test/Hab, GFP 0.71 ± 0.12, ChR2 0.37 ± 0.05, Unpaired T-Test p = 0.03) (Fig S7). As reduced activity is closely associated with defensive behavior in rodent models these results support the premise that stimulation activated amygdala circuits that encode for aversive signals.

In contrast, photo-activating the NBm-BL pathway led to ChR2 mice spending significantly more time than controls in the light-paired region (Conditioning: GFP (n = 8 mice), 41.1 ± 11.5%; ChR2 (n = 11 mice), 73.1 ± 8.4%; Two-Way ANOVA Interaction F(2,34) = 4.381 p = 0.02; Bonferonni post-hoc; GFP vs ChR2, Cond p = 0.03; Hab vs Cond, ChR2 p = 0.008; Cond vs Test p = 0.01) (Fig 5A,B). Compared to baseline conditions, during conditioning mice spent more time in the stimulation-paired region (Normalized data: Cond/Hab GFP = 0.96 ± 0.31, ChR2 = 1.67 ± 0.13, Unpaired T-Test p = 0.04) (Fig 5C-left) but not during the testing phase (Normalized data: Test/Hab GFP = 0.92 ± 0.22, ChR2 = 1.26 ± 0.16, Unpaired T-Test p = 0.23) (Fig 5C-right). To further probe the impact on NBm-BL inputs on BL reward circuits we implemented a self-stimulation paradigm where NBm-BL stimulation was paired with an illuminated touchscreen (Fig 5G-I). During testing, when mice only received stimulation when touching the paired touchscreen, ChR2 animals performed more correct touches to receive stimulation (Day 1: GFP, 5.5 ± 2.1; ChR2 7.4 ± 1.1; Day 2: GFP, 3.3 ± 0.6; ChR2, 14.5 ± 2.7; Two-way ANOVA Day, F(1,10) = 7.03 p = 0.024, Interaction Day*Group, F(1,10) = 5.5 p = 0.04; Bonferonni post-hoc Day 2 GFP vs ChR2, p = 0.002, ChR2 Day 1 vs Day 2, p = 0.006) (Fig 5H), supporting the idea that NBm-BL activation can reinforce appetitive instrumental learning. Increases in correct touches were unlikely due to increases in locomotion as beam breaks were similar between the groups (Day 2: GFP, 309.2 ± 42.3; ChR2, 376.3 ± 45.0; Unpaired T-Test t(10) = 0.93, p = 0.37) (Fig 5I).

**Figure 5:**
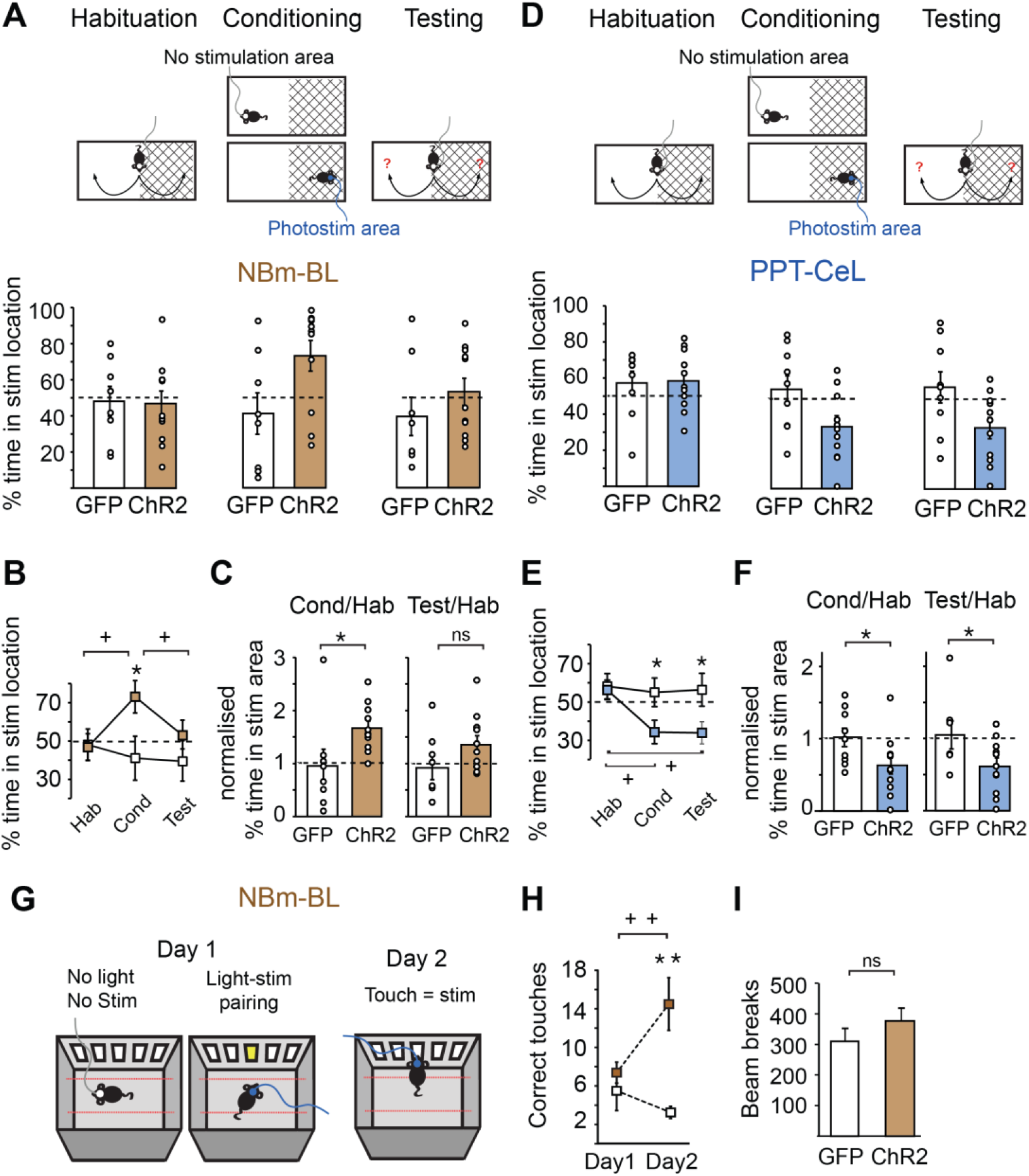
NBm to BL and PPT to CeL inputs, respectively, drive approach and avoidance behavior. A,D. (top) Schematic of the place preference paradigm where animals underwent a habituation, conditioning, and testing phase and raw data (circles) including averages (bar), of the time spent in the stimulation area for all phases in NBm-BL (A, bottom) and PPT-CeL (D, bottom) mice. In the place preference paradigm, ChR2-exprerssing mice receiving NBm-BL stimulation spent an equal amount of time in each region during the habituation phase but spent less time than control GFP-expressing mice in light-paired region during stimulation (B). For NBm-BL ChR2 mice, the ratio of stim time spent in conditioning phase over time spent in the habituation phase was significantly greater than GFP mice (C). Mice receiving PPT-CeL stimulation similarly showed no region-selective difference during the habituation phase but instead spent less time than control mice in only the light-paired region during the conditioning and testing phases (E). For PPT-CeL ChR2 mice, the ratio of stim time spent in both conditioning and testing phases over time spent in the habituation phase was significantly less than GFP mice (F). G. Schematic of the self-stimulation paradigm. On Day 1 mice received center screen light photo-stimulation pairings. On Day 2 mice received photo-stimulation only when the center touchscreen was touched. On Day 1 of a self-stimulation paradigm, ChR2 (n = 8) and GFP (n = 6) mice touched the paired touch screen an equal number of times while on Day 2 ChR2 mice correctly touched more than GFP-control mice. C3 (H). Locomotor activity as measured by beam breaks was no different in control and ChR2 groups (I).

In light of the above evidence demonstrating that NBm- and PPT-ChAT projections to the amygdala can convey information regarding valence, we next aimed to determine whether these ChAT neurons respond naturally to appetitive and aversive stimuli in wild-type mice (Fig 6). Here, retrograde-labeling fluorescent beads were injected in the BL or CeL of mice and each group was subject to a behavioral protocol where they either had access to a sucrose solution (appetitive) or exposed to unsignaled mild electrical footshocks (aversive). Mice were subsequently perfused and brains were processed for visualizing the protein product of the immediate gene cFos – a commonly used marker of neuronal activity – for revealing active BL- and CeL-projecting NBm- and PPT-ChAT neurons, respectively (Fig 6). Approximately two-thirds of retrogradely labeled NBm-BL and one-half of PPT-CeL neurons were immunopositive for ChAT (n = 8, 6 mice, 656, 395 cells: 64%, 45%, respectively) (Fig 6C,H). Notably, BL and CeL retrobead injections did not also label cells in the PPT and NBm, respectively, indicating that neither the NBm or PPT project to both BL and CeL. Compared to control mice that received unsweetened drinking water, sucrose-exposed mice had significantly more BL-projecting NBm-ChAT neurons express cFos (Ctl: n = 5 mice, 10.1 ± 2.1%; sucrose, n = 5 mice, 23.9 ± 5.4%; Unpaired T-Test, p = 0.04) (Fig 6E). In contrast, BL-projecting NBm-ChAT neurons in mice exposed to footshocks were not more active than control (n = 3 mice/condition: Ctl, 2.0 ± 1.1%; footshock, 0.0. ± 0.0%; Unpaired T-Test, p = 0.19) despite clear differences in defensive behavior (n = 3 mice: Ctl, 8.3 ± 1.1%; footshock, 62.7 ± 3.1%; Unpaired T-Test, p = 0.002) (Fig S7). Surprisingly, in mice exposed to shock few NBm neurons in total were immunopositive for cFos (n = 3 mice/condition: Ctl, 9.0 ± 3.0; footshock, 8.0 ± 2.4). Expanding on previous studies [28–31], these results demonstrate that a subpopulation of NBm neurons that respond to appetitive stimuli project to the BL. In CeL-injected mice, PPT-ChAT neurons projecting to the CeL in shock-exposed mice were not more active than in control mice (Ctl, 1.5 ± 0.9%; footshock, 1.0 ± 0.9%; n = 4 mice/group, Unpaired T-Test, p = 0.71) (Fig 6J). However, consistent with previous research [55], the scarcity of cFos expressing PPT-ChAT neurons makes this data difficult to interpret.

**Figure 6:**
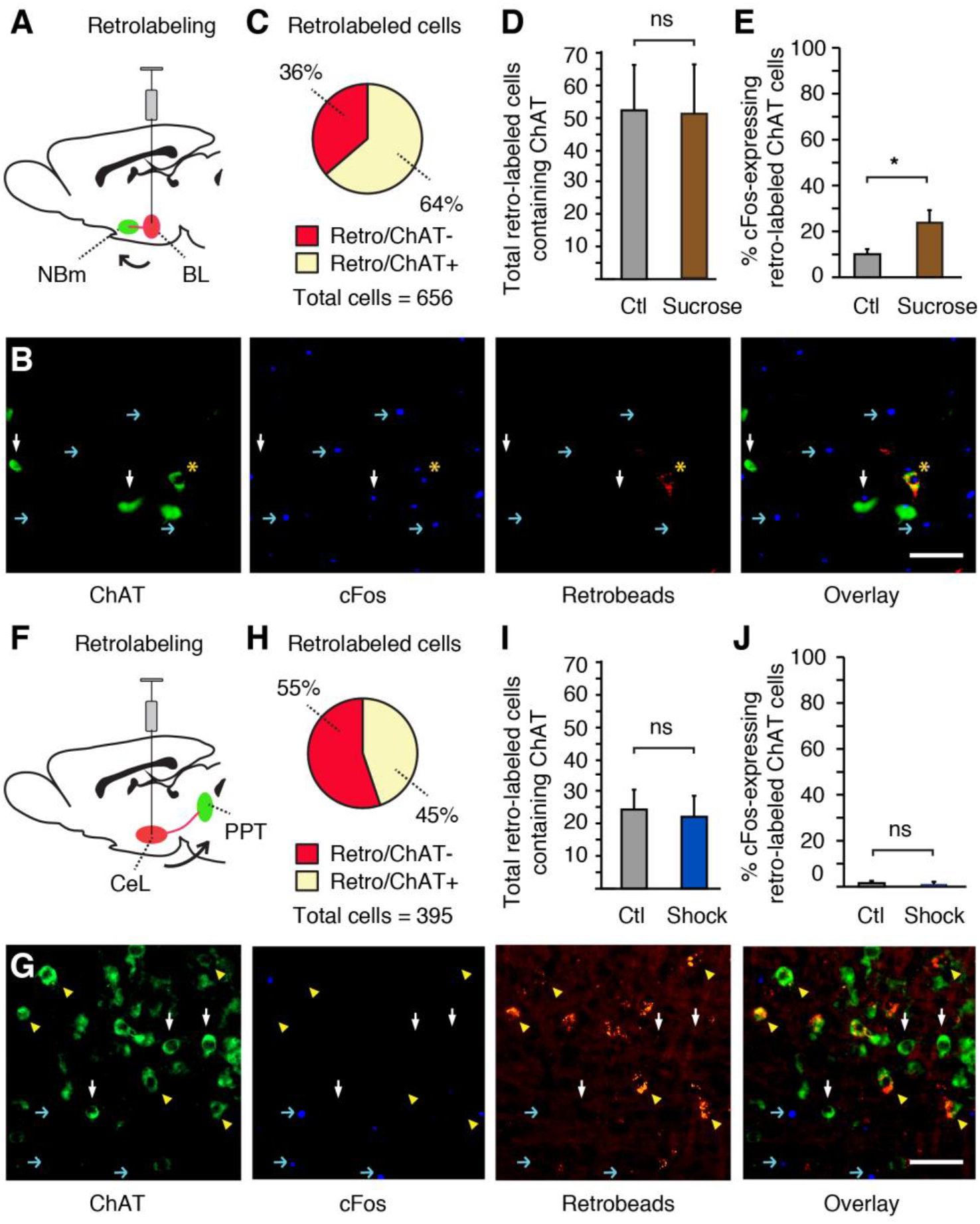
Appetitive stimulus activates BL-projecting NBm ChAT neurons. A. BL injections of retrobeads retro-labeled both ChAT+ and ChAT- NBm neurons (B,C). In control and sucrose-exposed mice, there was an equal amount of NBm-projecting ChAT+ neurons (B,D). Sucrose exposed mice however had a greater amount of retro-labeled NBm ChAT neurons express cFos than control mice (E). F. CeL injections of retrobeads retro-labeled both ChAT+ and ChAT- PPT neurons (G,H). Control and foot-shock exposed mice had similar amounts of CeL-projecting ChAT neurons in the PPT (G,I). Mice receiving footshocks showed no difference in the amount of retrolabeled PPT ChAT neurons that express cFos compared to control mice (J). White arrows, ChAT-expressing neurons only; blue arrow, cFos; yellow arrowheads, retrogradely-labeled ChAT expressing neurons; yellow star, retrogradely-labeled ChAT neurons expressing cFos. Scale bars: B, G, 50 μm.

## Discussion

Our results demonstrate that cholinergic projections to distinct amygdala nuclei can differentially impact amygdala circuits through unique modes of communication engaging both cellular and network-level processes that are known to support plasticity. Consistent with this, photo-activating NBm-BL and PPT-CeL pathways induced learning-related approach and avoidance behavior, respectively. These results reveal distinct cholinergic influences on amygdala circuits and motivational behavior.

### Principal amygdala neurons are synaptically influenced by both fast and slow cholinergic transmission

Central cholinergic transmission occurs through synaptic [56] and volume transmission [57]. Consistent with previous work [45], in the current study NBm-ChAT terminals reliably inhibited a high percentage of BL neurons via a M1 receptor-dependent increase in GIRK channel conductance. Both synaptic and extra-synaptic M1 receptors are expressed in the amygdala [58,59] indicating that the physostigmine-sensitive inhibitory responses measured were possibly due to the synaptically-released ACh acting at or near the synapse. In contrast, the slow M1-mediated excitation may arise from a separate class of extra-synaptic M1-receptors that is coupled to M1-mediated decreases in K^+^ conductance [60]. Our results provide support for both synaptic and volume ACh transmission via separate M1-mediated mechanisms for coordinating BL activity. Although the inhibitory response is likely due to ACh terminals in close proximity to BL principal neurons, is in unknown whether or not the relatively slower depolarizing effects are mediated via the same neurons or through nonspecific increases in ACh arising from the dense axonal arborization of cholinergic BL inputs.

### Rhythmic coordination of amygdala activity by NBm cholinergic populations

In many instances biphasic inhibition-excitation responses were evoked in BL principal neurons. When NBm-ChAT inputs to the BL were similarly stimulated *in vivo*, the rhythmicity of NBm and BL activity was significantly increased. The increased synchrony in theta frequency that outlasted NBm-ChAT stimulation was dependent on muscarinic receptor activation while nicotinic receptor activation contributed to NBm-evoked gamma oscillations in the BL. In addition to influencing principal neurons, NBm-ChAT projections to the BL have been shown to depolarize late-spiking (LTS) interneurons via nicotine receptors [45]. Through direct muscarinic inhibition-excitation and indirect nicotinic-receptor mediated inhibition, these results indicate that an interplay between both receptor types acting on different BL cell populations can effectively synchronize BL networks.

Cholinergic circuits promote neural oscillations in efferent regions [19–21] and behaviorally, oscillations in the theta and gamma frequency band have been linked to learned [43,44] and innate emotional responses [41,42]. Interestingly, in the latter case, coherent prefrontal-amygdala activity is strongly related with fear or safety signaling [41,42]. As the NBm can synchronize amygdala and prefrontal [61,62] activity our results raise the possibility that such behaviorally-relevant increases in PFC-amygdala coherence may in part stem from coordinated NBm activity that influences both structures.

### Brainstem ChAT neurons excite CeL amygdala circuits by synergistically activate AMPA and NMDA receptors through glutamate release

PPT-ChAT evoked CeL responses *in vivo* were insensitive to ACh antagonists. Interrogation of this circuit *ex vivo* revealed that these responses were instead due to glutamatergic excitation. Prior studies have demonstrated that PPT-ChAT neurons express glutamate markers [49] and that glutamate released from cholinergic neurons can effectively impact postsynaptic targets [50,56]. Despite PPT-ChAT neurons projecting to the CeL (Fig 6G,H), neither applied ACh nor the stimulation parameters used evoked a cholinergic receptor-dependent response on CeL neurons, suggesting that the predominant impact of PPT-ChAT terminals in the CeL is via the release of glutamate.

PPT-ChAT to CeL stimulation led to a sustained NMDA-dependent increase in excitability only at V_m_ close to threshold suggesting that PPT inputs are most effective at regulating CeL activity at depolarized levels. It is well established that by acting as activity-based coincidence detectors, NMDA receptors are critical for synaptic plasticity that underlies learning and memory. NMDA currents in CeL neurons have been shown to increase in amplitude following fear conditioning [63]while blocking them reduce both auditory and contextual fear conditioning [64], demonstrating a functional role for these receptors in fear learning. Interestingly, the response kinetics of CeL neurons are known to be relatively slow compared to those in BL principal neurons [65]. It is therefore conceivable that, by activating NMDA receptors on CeL neurons, PPT-CeL inputs would result in greater calcium entry and generate a longer time window for synaptic integration onto CeL neurons, thus providing a synaptic mechanism whereby PPT-ChAT inputs to the CeL can support plasticity.

### BL and CeL inputs from distinct ChAT cell populations drive opposing approach and avoidance learning-related changes in behaving mice

Our behavioral data indicates that activating NBm- or PPT-ChAT inputs to the BL and CeL can reinforce behavior by respectively biasing amygdala processing towards approach and avoidance behavior.

These results point towards a complementary role of distinct cholinergic influences on distinct amygdala circuits and functions. Our data suggests that PPT-CeL inputs preferentially contact CeL microcircuits important for fear learning instead of those important for appetitive emotional processes [66]. Approximately 70% of PPT-ChAT connected CeL neurons showed signature late-spiking properties while the rest were of the regular spiking variety. In seeming conflict with our data, late-spiking PKCδ^+^ CeL neurons are known to reduce the expression of defensive behavior [67]. Li et al., [63]however have shown that somatostatin (SOM)-expressing CeL neurons important for fear memory also show late-spiking and regular-spiking properties. As late-spiking neurons make up a third of CeL neurons [68] it is likely that multiple classes exist for differently regulating amygdala-dependent processes.

### Cholinergic inputs to the basolateral division on the amygdala drive reward-based learning

The behavioural results indicate that NBm-ChAT inputs to the BL support approach behavior. Moreover, NBm-ChAT neurons projecting to the BL were activated during reward delivery but not to aversive stimuli. These results conflict with a recent paper showing that activation of BL-projecting NBm-ChAT neurons can support fear learning [46]. This seeming contradiction may arise from the fact that in the Jiang et al. study NBm-BL axons were photo-activated during tone-shock pairings thereby influencing robust naturally occurring co-activated amygdala inputs. In the current study, NBm fibers were activated only when mice were exposed to neutral environmental stimuli.

Neurons within the BL respond to stimuli with positive and negative valence [37,38]. As such, it is hard to reconcile how such a dense ACh input can preferentially contact neurons that make up appetitive amygdala circuits. The rostral-caudal position alone cannot account for these findings since we positioned the fiber-optic implants where both positive and negative valence-coding neurons reside [69]. Our results can perhaps be partially accounted for in the functional division of the LA and BL. In the current study, the segregated NBm-ChAT projection to the BL allowed us to pinpoint the specific influences of ACh populations on BL function that is independent of the LA. Despite much experimental evidence demonstrating that LA and BL neurons respond to both positive and negative valence stimuli, data also exists showing that the LA and BL may in some instances be performing non-overlapping functions. For example, lesions restricted to the LA impair fear conditioning while BL selective lesions do not [70,71]. Moreover, LA neurons undergo fear-related plasticity [43,72] and activating LA populations can support fear memory [73] while BL neurons can also respond to safety signals [74] and can drive reward seeking [54]. Interestingly, recent work by Lee et al. [75]shows that increases in BL activity correspond more with the behavioral output of reward-seeking than to the stimulus driving the response, irrespective of the behavioral output being conditioned or spontaneous. This data suggests that specific BL circuits, rather than mostly communicating sensory-related information, are wired for generating appropriate motivation-based behavior, which is an idea consistent with our data suggesting that NBm-ChAT neurons convey value-based information to the BL.

Our results show that coordinating BL activity with NBm-ChAT activation can result in positive reinforcement. Moreover, BL-projecting ChAT neurons respond to appetitive but not aversive stimuli. Since NBm neurons rapidly respond to natural and learned emotionally-salient stimuli [28–31] it is possible that salient appetitive information is relayed to the amygdala through ACh neurons, thereby contributing to BL valence coding of rewarding stimuli that can ultimately support BL-dependent plasticity and learning and memory. In line with this idea, pairing an auditory stimulus with NBm-stimulation can result in associative memory [76] and recent work by Jiang et al. [46] has shown that stimulation of NBm-BL inputs *ex vivo* is sufficient to induce long-term potentiation of synaptic responses in the BL. Collectively, these results support the idea that by way of coordinating network activity, NBm-ChAT inputs to the BL may be providing the physiological substrate necessary for detection of appetitive stimuli, that are important for reward-based learning and memory.

In summary, our results reveal complementary anatomical, synaptic, network-level and behavioral cholinergic influences on amygdala microcircuits. As cholinergic neurons and amygdala function are critical for learning and memory our results demonstrate distinct mechanisms through which central ACh populations can engage processes known to promote associative biophysical changes that underlie memories. By doing so these results advance our understanding of the underlying neural mechanisms supporting emotional learning and consequently those affected in diseases such as Alzheimer’s Disease, where severe disruptions in central cholinergic systems result in disabling impairments in emotional memories.

## Acknowledgements

This work was supported by the Royal Society, The Wellcome Trust (JAS), the Sigrid Juselius Foundation (TA), and by a Herchel Smith Fellowship (YAH).

## Author Contribution

Conceptualization, T.A., Y.A.H., B.U.P., J.A-S.; Methodology, T.A., Y.A.H., B.U.P., J.A-S.; Formal Analysis, T.A., Y.A.H., B.U.P., J.A-S.; Investigation, T.A., Y.A.H., B.U.P., J.A-S.; Resources, L.M.S, T.J.B., O.P., J.A-S.; Writing – Original Draft, J.A-S. Writing – Review and Editing, T.A., Y.A.H., B.U.P., T.J.B., O.P., J.A-S.; Visualization, T.A., Y.A.H., J.A-S.; Supervision, J.A-S., Project Administration, J.A-S.; Funding Acquisition, L.M.S, T.J.B., O.P., J.A-S.

## STAR Methods

CONTACT FOR REAGENT AND RESOURCE SHARING

Further information and requests for resources and reagents should be directed to and will be fulfilled by the Lead Contact, John Apergis-Schoute

### EXPERIMENTAL MODEL AND SUBJECT DETAILS

Male heterozygous adult mice expressing Cre recombinase under the control of the ChAT promoter [ChAT-cre mice (B6N;129S6-Chattm2(cre)Lowl/J), The Jackson Laboratory, Bar Harbor, ME, USA] were used in the experiments. Mice were housed 2–10 animals per polycarbonate cage and provided ad libitum with water and standard lab chow (RM3, Special Diet Services, Essex, UK) in a holding room maintained under 12-h light cycle (lights off at 7 p.m.) with temperature regulated at 22–24°C and relative humidity kept at 50–55%. Mice were genotyped using PCR from ear notch biopsy. Before commencing behavioural testing the animals were handled daily for one week, and 1 h before each behavioural trial the mice were transported to the respective testing area. The experiments were performed during the light phase of the light/dark cycle. All animal care and experimental procedures were conducted in accordance with the UK Animals (Scientific Procedures) Act, 1986.

### METHOD DETAILS

#### Neurosurgery, Viral Transfection & Fiber Optic Implants

ChAT:cre mice at the age of 2.5 months were anaesthetised with isoflurane (5% induction, 1–2% maintenance, Abbott Ltd, Maidenhead, UK) mixed with oxygen as a carrier gas (flow rate 0.8–1.0 L/min), placed in a stereotactic frame (David Kopf Instruments, Tujunga, CA, USA), the skull exposed via a small incision, and a small bilateral craniotomy was performed to allow intracranial injections and placement of fiber optic implants. A stainless steel bevelled microinjector was lowered to a coordinate aimed at the basal forebrain (anteroposterior +0.05 mm in relation to bregma, laterally +/− 1.15 mm in relation to midline and at –5.0 mm from the skull level) and brain stem PPT (anteroposterior +4.60 mm in relation to bregma, laterally +/− 1.00 mm in relation to midline and at –3.50 mm from the skull level). Microinjector was connected to a 1 μL Hamilton glass syringe via polyethylene tubing and injection rate of 0.1 μL/min was regulated by a microprocessor controlled programmable syringe pump (KD Scientific Inc., Holliston, MA, USA). Injection site received 200 nL volume of one of the following viruses: AAV5-EF1α-DIO-GFP or AAV5-EF1α-DIO-hChR2(H134R)-GFP, (titer 4x10^12^ vg/mL, University of North Carolina, Gene Therapy Center, NC, USA) followed by 5-min wait. For postoperative care mice received meloxicam (1 mg/kg. s.c., Boehringer Ingelheim Ltd. Bracknell, UK) and a recovery period of 4 weeks was attained before commencing fiber optic implantation.

For behavioural experiments, chronic fiber optic implants were fabricated by connecting stripped optic fiber (200 μm, FT200EMT 0.39NA, Thorlabs, Newton, NJ, USA) to ferrules (CFLC230,Thorlabs) as reported previously (Sparta et al. 2012, Nat Protocols 7:12–23). Using the same inhalation anaesthesia protocol as for viral transduction and surgical preparation, the skull was exposed and fiber optic implants were lowered to a coordinate at the basolateral amygdala function amygdala for the basal forebrain injected animals (anteroposterior –1.50 mm in relation to bregma, laterally +/− 2.70 mm in relation to midline and at –4.00 mm from the skull level) or central lateral amygdala for the PPT injected animals (anteroposterior –1.20 mm in relation to bregma, laterally +/− 2.30 mm in relation to midline and at –4.40 mm from the skull level). Fiber optic implants were attached to the skull using two miniature screws and dental cement. Meloxicam (2 mg/kg s.c.) was administered as analgesia and animals were let recover in a postoperational chamber with maintained +36.0°C.

#### Electrophysiology and in vitro photostimulation

Coronal slices were made >9 weeks post-injection as in our previous study. 250 mm thick slices were cut with a Leica VT 1200S vibratome in ice-cold ACSF (see below), and allowed to recover for 1 hour at 35 °C in ACSF before recordings. Patch pipettes were manufactured from borosilicate glass, and their tip resistances were 4–6 MΩ when filled with K-gluconate solution (see below). Whole-cell recordings were carried out at 37 °C using an EPC-10 amplifier and PatchMaster software (HEKA Elektronik, Germany). Only cells with access resistances of <20 MΩ were used for analysis. Current signals were low-pass filtered at 3 kHz and digitized at 10 kHz. Data were analyzed with Minianalysis (Synaptosoft), Igor and Adobe Illustrator/ Photoshop. Whole-cell recordings were performed at 35°C using an EPC-10 amplifier and Patch-Master software (HEKA Elektronik). ChAT-containing cells were visualized in acute living brain slices using a GFP filter set (Chroma). To stimulate ChR2, we used a ThorLabs blue LED, delivering ∼10 mW/mm2 light to ChR2-containing axons around the recorded cell 40× 0.8NA objective. Averaged data are presented as the mean ± SEM. Slice-cutting and recording ACSF was gassed with 95% O2 and 5% CO2, and contained the following (mM): NaCl 125, NaHCO_3_ 25, KCl 3, NaH2PO4 1.25, CaCl2 1/2 (cutting/recording), MgCl2 6/1 (cutting/recording), sodium pyruvate 3 and glucose 25/5 (cutting/recording). Pipettes were filled with (in mM): potassium gluconate 135, NaCl 7, Hepes 10, Na2-ATP 2, Na-GTP 0.3, and MgCl2 2; pH was adjusted to 7.3 with KOH. All chemicals were from Sigma, Tocris, and Abcam.

#### Behavioral Assays

##### Animals

All animals were initially weighed for 3 consecutive days to establish free-fed weights. Weights were then restricted to approximately 85% of their free-fed weight via daily provision of standard laboratory chow. Water was available ab libitum throughout the study. Animals were kept under a 24h light/dark cycle (lights on 0700, lights off 1900). All behavioral testing was carried out under the light cycle. Prior to behavioral testing, animals were removed from the home-room and taken to the testing room. Following completion of daily testing, animals were returned to the home-room.

##### Place conditioning

Behavioral tests were started by habituating animals to the connecting of laser patch cables with the fiber optic implants. The conditioning apparatus consisted of a chamber (45 × 23 × 20 cm) with two distinct equal sized contextual zones that differed in their tactile floor surface (mesh and smooth). There was no wall or door between the zones and animals were thus free to move between the contextual spaces. The location and movement data were collected by video tracking using Ethovision XT software (Noldus Information Technology bv, Wageningen, The Netherlands). On day one, in a habituation test, animals were connected to the laser patch cables and let freely explore the chamber without laser stimulation. On day two in a 15-min conditioning trial, laser stimulation was delivered every time the animal was in the stimulation-paired zone, and the stimulation was not delivered when the animal was in the non-stimulation zone. On the day three in a 15-min testing trial, the animals were again connect to the laser, placed in the chamber with the floors, and let freely move in the chamber without laser stimulation. The laser stimulation-floor type pairing and the orientation (left-right) of the floor materials in the chamber were counterbalanced within the experimental groups.

##### Self-Stimulation

The Bussey-Saksida touchscreen chamber, a trapezoidal operant chamber, is contained within a sound-attenuating box. One end of the chamber is comprised of a touchscreen. Reponses are registered by breaking a set of infra-red beams located less than 5mm from the surface of the screen. At the other end of the chamber a small aperture standardly delivers liquid reinforcer. This was blocked by a metal plate throughout the duration of this study. Throughout testing, a standard ‘5-choice’ Perspex mask, which comprises 5 evenly spaced apertures, was placed in front of the touchscreen to focus responses and protect the edges of the screen. Two sets of infrared beams (at the front and back of the box) were used for measurement of locomotor activity and initiation of task events.

On the first day of testing, all animals were placed on an ‘initial touch’ schedule to establish an association between the central stimulus and stimulation. Under this schedule, the central location of the 5-choice mask was illuminated by a white square stimulus for 30 seconds. Following this, the illumination ceased and the animal immediately received stimulation for 10s. If the illuminated location was touched during the 30s window, the illumination immediately ceased and the animal received 30s of stimulation. A front beam break was required to initiate the next trial. These sessions terminated after either 30 successfully completed trials or 60 minutes. On the second day of testing, stimulation was controlled by a fixed ratio 1 schedule (FR1), which required animals to make a single touch at the center location for 10s of stimulation. Illumination at the central stimulus ceased when the stimulus was touched. A new trial was initiated by breaking the front beam. The center location was illuminated until a successful center touch was made. This session was terminated following either completion of 30 trials or after 60 minutes had elapsed. Following completion of behavioural tests, measures for trials completed, blank touches, and front and back beam breaks were extracted from Abet Touch II testing software and analyzed.

#### Optical stimulation in vivo

Laser light was generated by using 473-nm diode laser (Versalase, Vortran Laser Technology Inc., Sacramento, CA, USA) controlled by Arduino Uno microcontroller (Arduino Community) to generate pulsing (20 Hz, 5 ms pulse length) with ∼15 mW laser power at the tip of the fiber optic implant. Prior to surgical implantation the implants were tested and accepted for implantation with laser light transmittance rate of 75–85% of power measured from patch cable output.

#### Acute in vivo recordings

##### Surgery and Light Delivery

Adult mice ChAT-Cre*Ai32 or ChAT-CRE injected with AAV5-EF1a-ChR2-eYFP at least 8 weeks prior to recordings. Mice were deeply anesthetized via intraperitoneal injections of urethane (1.5 to 1.8 g/kg injected in several shots until loss of reflexes is assessed by paw pinching). While restrained in a stereotaxic apparatus (David Kopf instruments, Phymep, France) their temperature was monitored and maintained at 34–36° C using a heating blanket (Basi). After exposing the skull, two small burr holes were drilled at NBm coordinates AP = +0.3 mm and ML = 3.0 mm and BA coordinates AP = -1.58 mm and ML = 1.2 mm relative to bregma.

Simultaneous optical stimulation and electrical recording in the NBm were carried out using an optrode which consists of an extracellular parylene-C insulated tungsten microelectrode (127-μm diameter, 1 MΩ; A-M Systems) tightly bundled with an optical fiber (200 μm, 0.22 N.A.; Thorlabs), with the tip of the electrode protruding 0.2–0.7 mm beyond the fiber end to ensure illumination of the recorded neurons as previously described (Shipton et al., 2014). Electrical recording from the amygdala were carried out using a simple extracellular parylene-C insulated tungsten microelectrode (127-μm diameter, 1 MΩ). The optrode was gradually descended into the NBm with a 20° angle. Response to photostimulation was regularly assessed between DV = 3.7 and DV = 4.1 mm to find the best location for an effective stimulation. A common ground electrode for the two electrodes was placed above the right part of the cerebellum and was submersed by saline solution (0.9% NaCl). The recording electrode was descended into the amygdala with a 15° angle to reach a final depth of DV=4.0mm. Light delivery was performed by collimating the optic fibre to a blue laser light (488nm, 15 +/− 3 mW at fiber tip; laser 2000). Light delivery was controlled by a shutter whose aperture was driven by custom-made Igor 6.5 procedures. Electrical signals were amplified and filtered at 0.1–5 kHz (1800 Microelectrode AC Amplifier; A-M Systems), digitized at 20 kHz using an ITC-18 board (Instrutech, Port Washington, NY) and Igor software (Wavemetrics, Lake Oswego, OR). All data were analyzed off-line using Igor Pro 6.37.

##### Freely moving recordings

Mice were handled for 2–3 weeks before the start of recordings to limit the amount of stress generated by the plugging of the pre-amplifier and optic fiber. For recordings, mice were placed in a large cage containing soaked pellet of food and bedding and were allowed to explore, eat and rest. Stimulation, recordings and analysis were performed using similar light source and control software as for anesthetized recording, except that the amplification and digitalization was performed using an in-built amplifier (Whisper system, Janelia, Research Campus, https://www.janelia.org/open-science/whisper).

##### Electrophysiology and Analysis

LFP signal was isolated from high-frequency signals using a low-pass filter cutting frequencies above 120 Hz and the signal was downsampled from 20 kHz to 1 kHz. Each stimulation procedure was repeated at least 20 times with a minimum of 20 s interval between each stimulation. Each recording was submitted to wavelet analysis and the power spectrum over time obtained by the wavelet analysis was z-scored for each frequency over the 20 s of the sweep. (z-score formula = fFreq(t) = (t-meanfreq)/sdFreq. The mean power spectrum for baseline and stimulation conditions was obtained by averaging the power spectrum over 0.5s preceding the stimulation and following the end of stimulation, respectively. Differences of the baseline and stimulation power spectrum were computed using 2-ways ANOVA. For these experiments, 5 ChAT-CRE * Ai32 and 2 transfected ChAT-CRE were used. The variation of the power spectrum between baseline and stimulation conditions was not significatively different between the 2 two pools of animals (2 ways ANOVA, for Amygdala p = 0.09864 and for SI p = 0.418159).

#### Immunocytochemistry

To determine the specificity of the opsin expression in the cholinergic neurons, the mice were anaesthesised with pentobarbital (500 mg/kg, i.p., Vetoquinol, Buckingham, UK) and transcardially perfused first with 0.1 M PBS followed by 10% neutral-buffered formalin (Sigma, St. Louis, MO, USA). Brains were removed, postfixed overnight at +4°C and cryoprotected at +4°C with 30% w/v sucrose in PBS until the brains sunk. Coronal sections (30 μm) were cut on a freezing sliding microtome (model 860, Americal Optical Company, Buffalo, NY, USA). For both GFP and VAchT immunostaining, the sections were washed at RT in 0.1 M PBS, blocked with 1% BSA (Fisher Scientific, Fair Lawn, NJ, USA) supplemented with 0.3% Triton X-100 (Fisher) in 0.1 M PBS. Sections were then incubated overnight at RT in primary antibodies diluted in blocking buffer, washed in PBS, incubated in secondary antibodies for 2 h at RT, washed in PBS, and mounted on microscope slides and coverslipped. Primary antibodies were chicken anti-GFP (1:1000, ab13970, Abcam, Cambridge, UK) and guinea-pig anti-VAchT (1:500, AB1588, Millipore, Temecula, CA, USA). Secondary antibodies were goat anti-chicken AlexaFluor 488 (1:1000, Abcam) and goat anti-guinea-pig AlexaFluor 594 (1:1000, Abcam). Digital images were captured with a Zeiss Axioskop 2 microscope (Zeiss, Oberkochen, Germany) and QImaging QICAM Fast digital camera (QImaging, Surrey, BC, Canada). Images were merged using ImageJ (National Institutes of Health).

#### cFos imaging of neuronal activation

##### Retrobead injections

C57Bl/6J mice at the age of 2.5 months were anaesthetised with isoflurane as described previously for viral transduction. A stainless steel bevelled microinjector was lowered to a coordinate aimed at the basolateral amygdala (anteroposterior –1.55 mm in relation to bregma, laterally +/− 2.70 mm in relation to midline and at –4.90 mm from the skull level) and the central nucleus of amygdala (anteroposterior –1.40 mm in relation to bregma, laterally +/− 2.30 mm in relation to midline and at –4.50 mm from the skull level). Injections were performed by using the same surgical injection system as with the viral transduction. Injection site received 200 nL volume of retrobeads (Lumafluor, Nashville, TN, USA) with a injection rate of 0.1 μL/min followed by a 5-min wait. For postoperative care mice received carprofen (5 mg/kg. s.c., Norbrook, Newry, UK) and a recovery period of 1 week was attained before the behavioural testing.

##### Sucrose delivery

After recovery mice were habituated to the operant chambers residing in sound-attenuating boxes (Med Associates, Fairfax, USA, operated by K-Limbic software, Conclusive Solutions, Sawbridgeworth, UK) for 3 consecutive days. For the sucrose delivery, the chambers were equipped with a food receptacle consisting a photobeam for detection of mouse nose-pokes. A nose-poke triggered a random (interval 3–17 s.) latency for delivery of a liquid sucrose reinforcer (15% w/v sucrose solution in water). Mice were given 90 min to consume sucrose, while for the control animals the reinforcer was not delivered. Immediately after the trial the mice were perfusion fixed as described above.

##### Foot shock delivery

After recovery mice were habituated to the fear conditioning chambers residing in sound-attenuating boxes (Med Associates, Fairfax, USA) for 3 consecutive days. For the foot shock delivery, mice received foot shock (1.2 mA, 2 s) on a pseudorandom interval (range 1–7 min) over a 30 min trial. For the control animals no foot shocks were delivered. The mice were returned to their home cages and 60 min afterwards perfusion fixed as described above. cFos immunohistochemistry:

##### cFos immunohistochemistry

To determine the neuronal activation of the retrogradely labelled ChAT-positive cells, triple-labeling cellular imaging was performed for retrobeads, ChAT and cFos. Brain samples were prepared for sections as described above. The sections were washed at RT in 0.1 M PBS, treated for 15 min with 10 mM citrate and for 1 h with hydrogen peroxide, blocked with 3% BSA (Fisher Scientific, Fair Lawn, NJ, USA) supplemented with 0.3% Triton X-100 (Fisher) in 0.1 M PBS. Sections were then incubated overnight at 4°C in cFos primary antibody diluted in blocking buffer, washed in PBS, incubated in secondary antibody for 2 h at RT followed by DAB intensification and washed in PBS. Sections were then incubated in a blocking buffer with 1% BSA supplemented with 0.3% Triton, and incubated overnight at 4°C with a ChAT primary antibody diluted in a blocking buffer, washed in PBS, incubated in secondary antibody for 2 h at RT, and mounted on microscope slides and coverslipped. Primary antibodies were rabbit anti-cFos (1:500, sc-52, Santa Cruz Biotechnology, Santa Cruz, USA) and goat anti-ChAT (1:100, AB144, Millipore, Temecula, CA, USA). Secondary antibodies were biotinylated horse anti-rabbit (1:200, BA-1100, Vector Laboratories, Burlingame, USA) and donkey anti-goat AlexaFluor 488 (1:1000, A-11055, Life Technologies, Eugene, USA). Digital images were captured with a Zeiss Axioimager microscope (Zeiss, Oberkochen, Germany) and Hamamatsu Orca Flash 4.0 digital camera (Hamamatsu, Shizuoka, Japan). Images were merged using ImageJ (National Institutes of Health).

### QUANTIFICATION AND STATISTICAL ANALYSIS

#### Statistical Analysis

Averaged data are presented as the mean ± SEM and all data was analyzed using two-tailed statistics. Each animal provided one value with repeated trials (i.e. stimulations) being averaged and the average providing the value for that animal. For cell counts, each n is data from one animal and for cellular ephys each n comes from a single cellular recording. Multiple cells were recorded from an individual animal but at least 3 mice were used for each in vitro dataset.

### DATA AND SOFTWARE AVAILABILITY

**Data will be available on request from corresponding author**.

---

**Supp 1:**
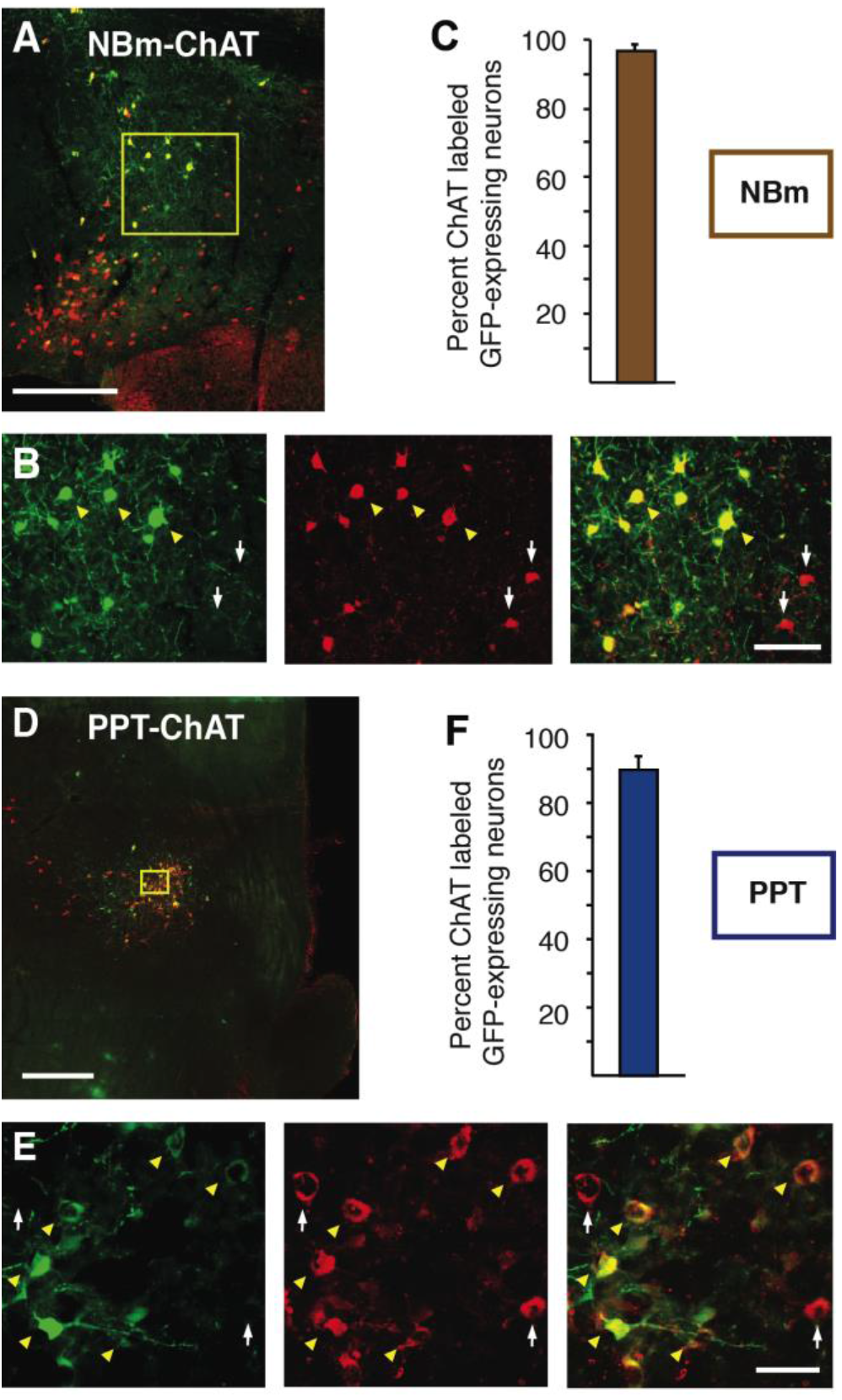
Viral-expression of GFP in ChAT-cre mice is highly specific to ChAT-containing neurons. Low (A) and high (B) magnification images of basal forebrain cholinergic population. Virally-expressed GFP (B; left,right) in NBm ChAT-containing (B; middle, right) neurons. D. PPT viral-injections labeled PPT ChAT-containing neurons with GFP (E; left, right) co-labeled with ChAT (E; middle, right). C,F. Summary data of GFP expression in NBm (C) and PPT (F) ChAT-immunoreactive neurons. White arrows, ChAT-expressing neurons only; yellow arrowheads, GFP and ChAT expressing neurons. Scale bars: A, 200 μm; B, 60 μm; D, 500 μm, E, 30 μm.

**Supp 2:**
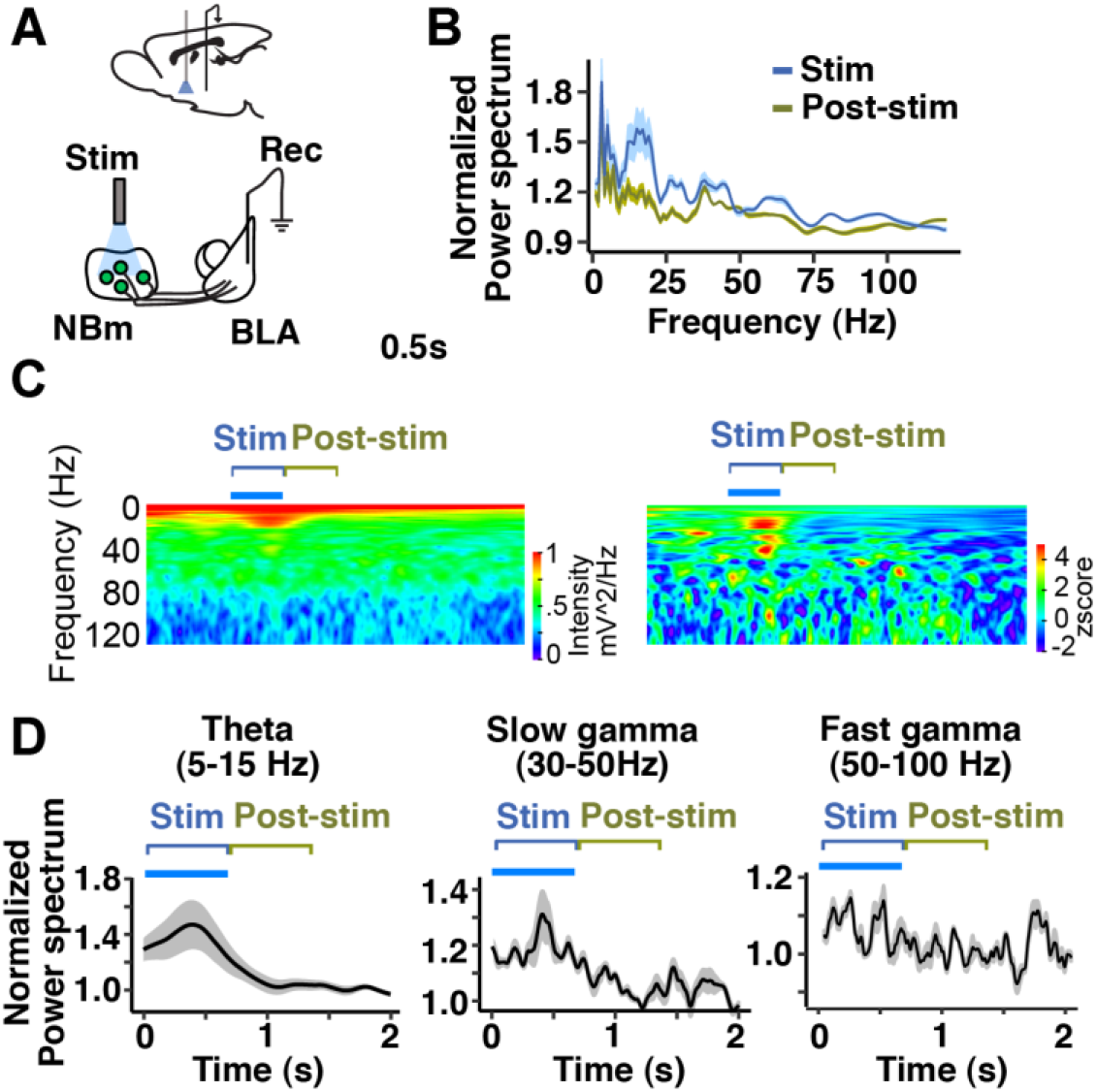
In vivo PPT/NBm photo-activation in awake mice synchronizes BL activity transiently in the theta and gamma frequency band. In freely-behaving mice NBm-ACh stimulation (A) results in an increase in theta and gamma power (B during and after stimulation (C,D). Power Spectrum units, mV^2^/Hz (C).

**Supp 3:**
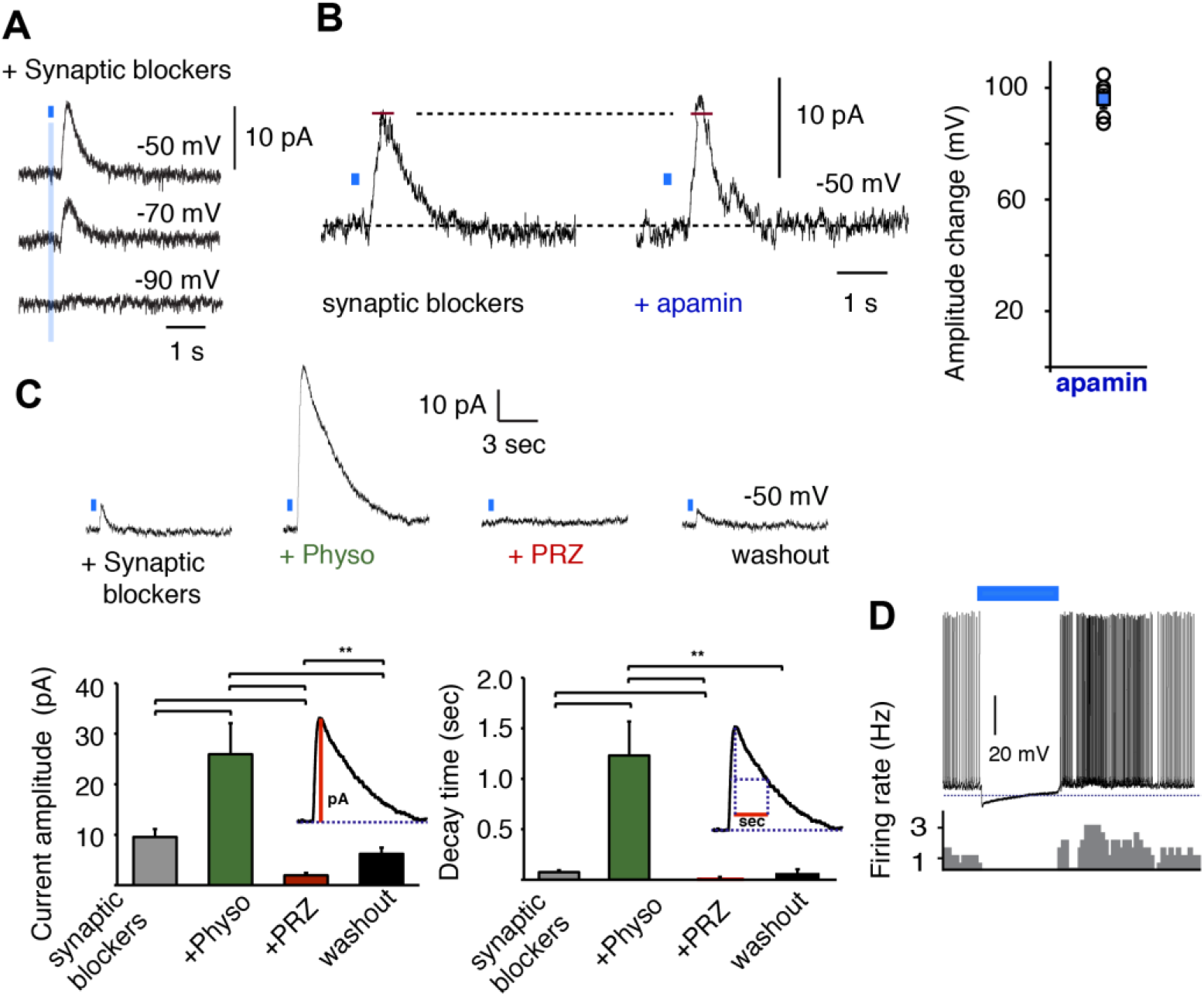
Synaptic properties of NBm inputs onto BL principal neurons. A. inhibitory currents evoked by minimal photo-activation of NBm terminals (0.5 ms flash) (n = 6) in the BL appeared 43.3 ± 2.5 ms after stimulation onset and reversed at approximately –90 mV. These inhibitory currents were unaffected by apamin, the SK Ca^2+^-activated K^+^ channel blocker (200 nM) (B) but increased in amplitude and duration following application of the cholinesterase inhibitor Physo (10 μM) (C). These responses were subsequently blocked by bath-application of the M1 receptor antagonist PRZ (C). D. In most cases 20 Hz stimulation for 20 seconds resulting in a clear biphasic inhibitory-excitatory response.

**Supp 4:**
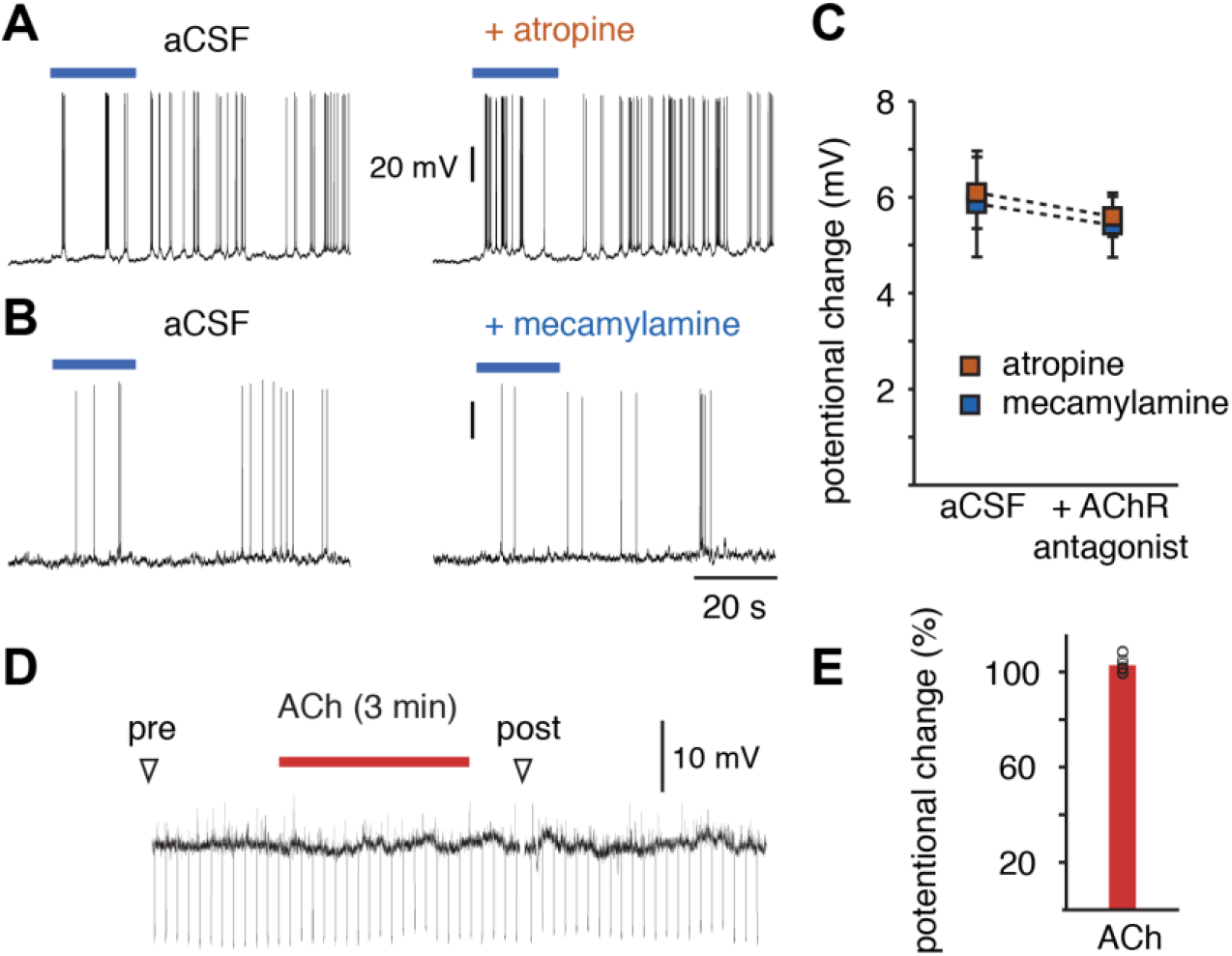
PPT-mediated excitation of CeL neurons is independent of ACh receptor activation. 20 Hz activation of PPT-CeL inputs is unaffected by ACh muscarinic (A,C) and nicotinic (B,C) receptor antagonists (ATR, 2 μM; MEC, 10 μM). D,E. PPT-CeL connected neurons were unresponsive to bath-application of ACh (100 μM).

**Supp 5:**
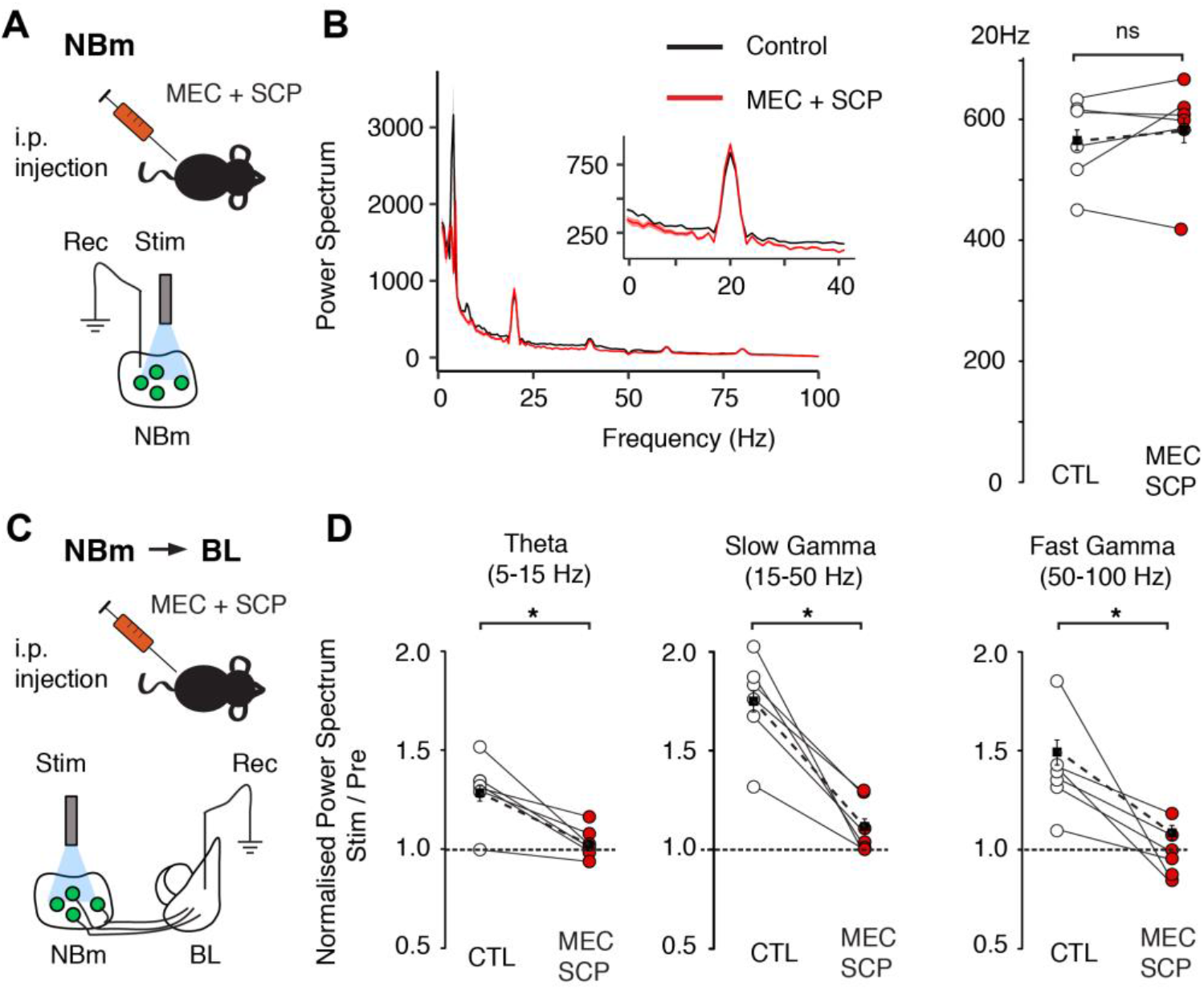
Intraperitonial injection of ACh antagonists diminishes NBm-mediated increase in BL network rhythmicity. A,B. ACh receptor antagonism does not significantly affect the local increase in 20 Hz power in response to 20 Hz photo-stimulation of NBm ChAT-containing neurons. C,D. In contrast, BL responses in different frequency bands (theta, slow and fast gamma) evoked by these same stimulations were significantly reduced by ACh receptor antagonism. Power Spectrum units, (B) μV^2^/Hz.

**Supp 6:**
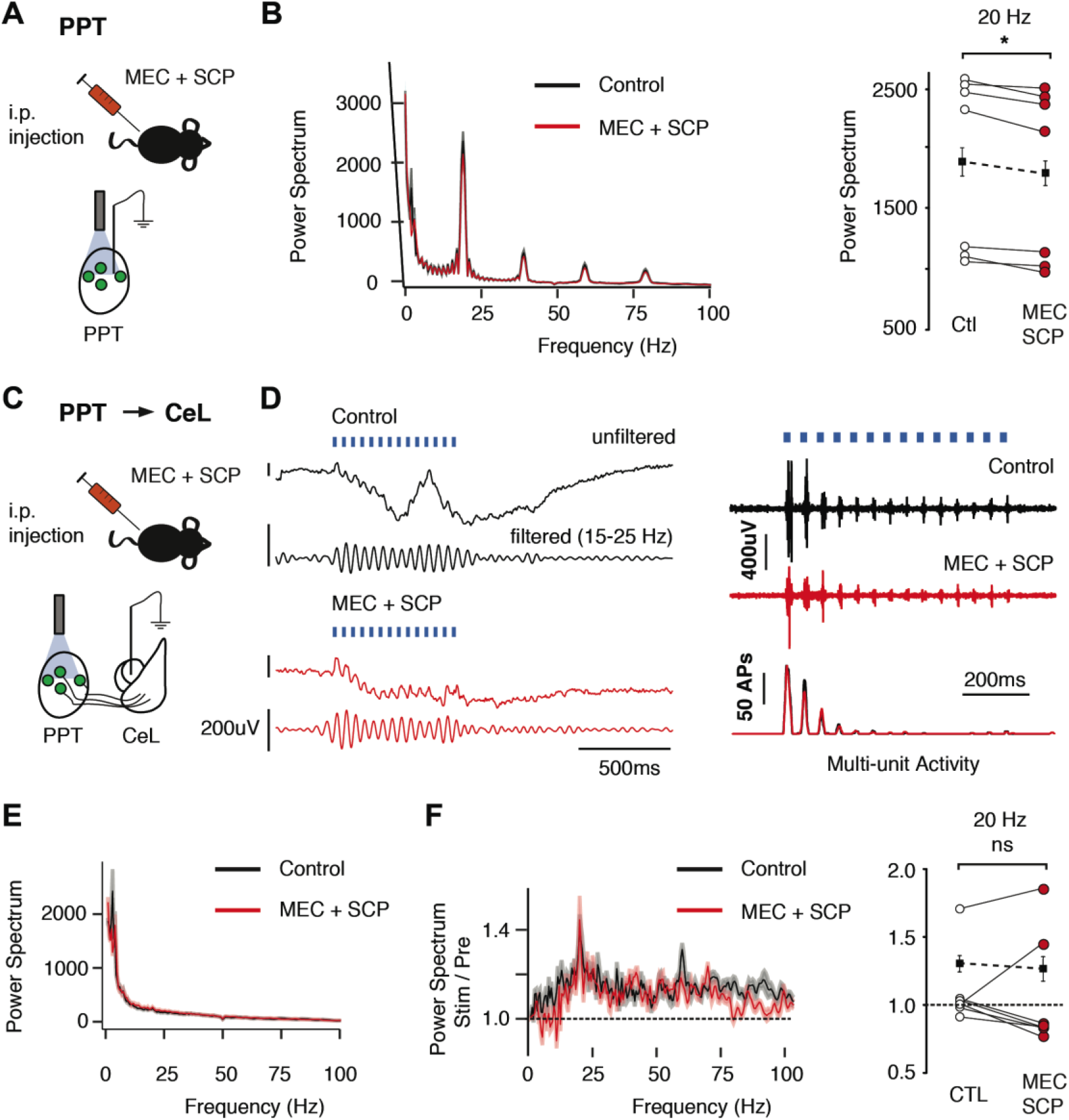
Intraperitonial injection of ACh antagonists diminishes PPT-mediated increase in local activity but has no effect on CeL networks. ACh receptor antagonism (A) significantly reduces the increase in 20 Hz power in response to 20 Hz photostimualtion of PPT ChAT-containing neurons (B-RIGHT). D. PPT-ChAT evoked field (D-left) and multiunit responses (D-right) confined to the CeL. Evoked field potentials were unaffected by ACh receptor antagonism (E,F). Power Spectrum units (B,E), μV^2^/Hz.

**Supp 7:**
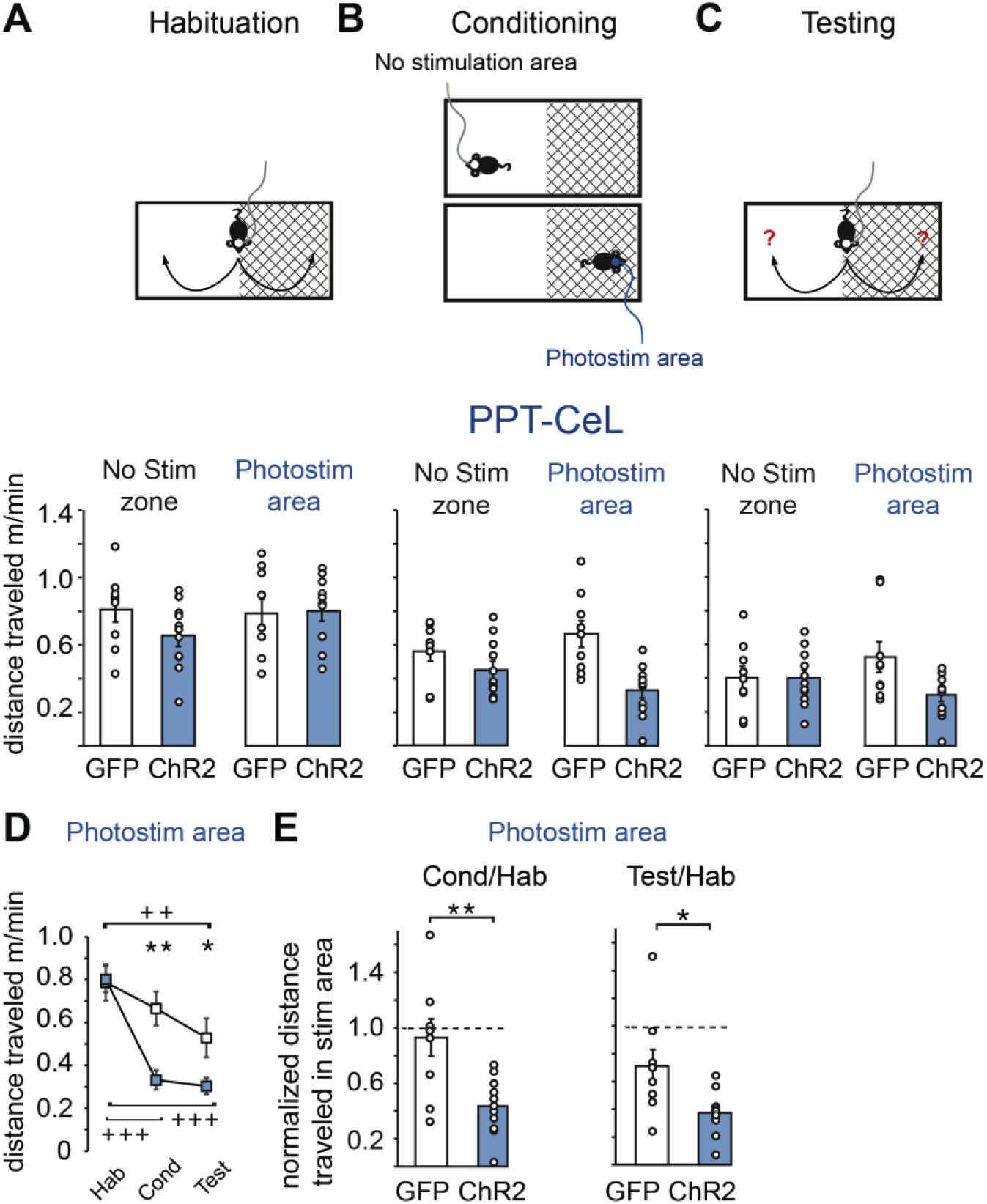
PPT-CeL activated mice were less active during place preference conditioning. A-C. Schematic of the place preference paradigm where animals underwent a habituation, conditioning, and testing phase (top) and raw data (circles) including averages (bar), of exploration speed (m/sec) for all phases in PPT-CeL mice (bottom). When in the stimulated area, PPT-CeL activated mice showed a significant reduction in speed during both conditioning and testing phases compared to the habituation phase and to GFP control mice (D). In the stimulation area, the rate of movement of ChR2 animals in both conditioning and testing phases normalized by their speed during the habituation phase was significantly less than GFP mice (E).

**Supp 8:**
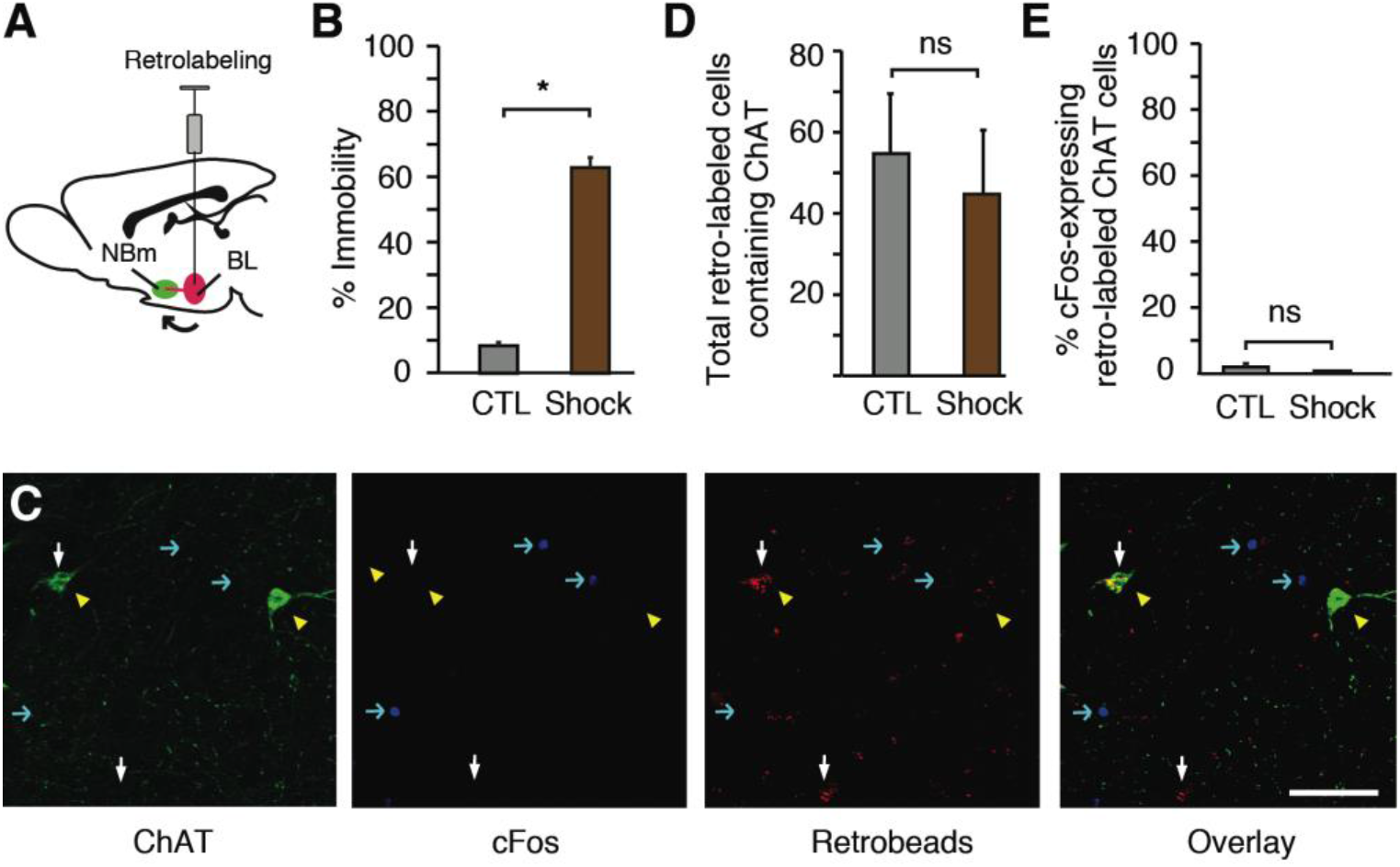
Aversive stimulus does not activate BL-projecting NBm ChAT neurons. BL injected mice with reotrobeads (A) expressed fear-related behavior (immobility) when presented with unpredicted footshocks (B). BL injections of retrobreads labeled NBm-ChAT neurons equally in control and footshock-exposed mice (C,D). No difference in cFos expression was seen between the groups (E). Scale bars: C, 50 μm.

